# Parental care is a genetic capacitor

**DOI:** 10.64898/2026.06.09.731115

**Authors:** Elise Parey, Thomas M. Houslay, Syuan-Jyun Sun, Angela T. Trowsdale, Daria Gavriouchkina, Modesta Blunskyte-Hendley, Rebecca M. Kilner, Ferdinand Marletaz, Rahia Mashoodh

## Abstract

Parental care is widespread across the animal kingdom and plays a critical role in offspring development. Yet, its broader genetic and evolutionary impacts remain underexplored. Here, using the biparental burying beetle *Nicrophorus vespilloides* as a model, we show that parental care acts as a genetic capacitor: it allows genetic variation to accumulate while care is present and releases it when care is disrupted. By experimentally manipulating care, we demonstrate that parental care suppresses genetic variation associated with offspring body size, which is released when care is lost. To investigate the underlying molecular mechanisms, we generate a chromosome-scale genome assembly for *N. vespilloides*, alongside a single-nucleus gene expression atlas and epigenomic datasets from larvae reared with and without parental care. We find that the loss of parental care induces molecular stress, disrupting the expression of the protein chaperone Hsp83, which is a well-known molecular capacitor, alongside other putative mRNA chaperones. Moreover, our results suggest that parental care buffers development by maintaining an open, responsive chromatin landscape and redundant gene regulatory interactions. Overall, our work reveals that parental care shapes the storage, expression and release of genetic variation with broad implications for adaptation and evolution.

## Introduction

Parental care is a major life-history adaptation that has evolved repeatedly across diverse animal lineages. While widespread in vertebrates including fishes, amphibians, birds and mammals, it has also recurrently evolved within various invertebrate groups, including arachnids, crustaceans and insects [1]. Through parental care, parents invest energy and resources to meet the demands of their developing offspring and improve their survival under challenging conditions. This investment encompasses shielding young from environmental stressors, ensuring nutrient access, reducing pathogen exposure, and constructing protective nests or microenvironments that support offspring development [1]. The biparental subsocial burying beetle, *Nicrophorus vespilloides,* has emerged as an ideal model to investigate the evolutionary and developmental consequences of parental care [2]. These insects are readily bred in the laboratory in large numbers across multiple generations. Crucially, care is facultative in this species, meaning offspring can self-feed and survive in its absence, making it possible to manipulate care experimentally and examine the genetic and regulatory programmes that respond to care [3].

One of the key functions of parental care is to stabilise or buffer the developmental environment of developing offspring. This model implies three key consequences. First, parental care directly improves offspring fitness by compensating for genetic or environmental vulnerabilities and, as a result, suppresses the expression of phenotypic variation despite underlying genetic diversity. Second, in doing so, care relaxes selection pressures allowing genetic variation to accumulate over generations, including deleterious mutations that would otherwise be purged. Third, changes in care or its loss could release this stored variation and expose it to selection, with potential consequences for adaptation that depend on the composition of the genetic variants accrued. Support for these predictions has been observed across diverse taxa. The fitness consequences of care, in particular, have been well demonstrated. For example, in dung beetles, the phenotypic effect of mutations is magnified under low care as opposed to high care [4]. Moreover, in both biparentally caring fish and burying beetles, inbred offspring suffer smaller fitness reductions when care is present [5–7]. Taken together, these observations raise the intriguing possibility that parental care may act as a genetic capacitor: storing heritable variation, buffering it from selection while care is present, and releasing it when care is disrupted.

However, definitive evidence supporting this hypothesis is lacking and the underlying molecular mechanisms remain undeciphered. Previous studies of genetic capacitors have focussed almost exclusively on the chaperone protein, Hsp83, as a candidate for developmental buffering [8–11]. Despite its name, Hsp83 is constitutively expressed during normal development rather than exclusively under stress [12]. It maintains protein homeostasis by refolding misfolded proteins that are a product of genetic mutations and interacts with various regulatory proteins and co-chaperons involved in developmental and genomic stability [12,13]. In studies of both *Drosophila* and *Tribolium*, the suppression of Hsp83 has been shown consistently to release cryptic genetic variation associated with morphological traits such as wings, eyes and legs, some of which can be adaptive [8–11]. As such, Hsp83 constitutes a relevant candidate for mediating parental care-driven developmental buffering, though this phenomenon is likely to emerge from the action of a number of interacting proteins, gene regulatory and signalling pathways [13,14].

Here, we use the burying beetle *N. vespilloides* to explicitly test the genetic capacitor model of parental care and investigate its underlying molecular mechanisms. We manipulated care to determine whether parental care effectively suppressed genetic variation in offspring body size. At the molecular level, we hypothesised that parental care may buffer development through the expression of Hsp83 and potentially other molecular capacitors that enhance proteostasis, chromatin stability and regulatory interactions between transcription factors, cis-regulatory elements and target genes. To address these questions and draw a comprehensive picture of molecular changes associated with the loss of parental care during early development, we generated a chromosome-scale genome assembly for *N. vespilloides* alongside omics datasets for the first-instar larval head. This includes a single-nucleus gene expression atlas (snRNA) and genome-wide chromatin accessibility (ATAC-seq) and regulatory activity (H3K4me3 Cut&Tag) profiling, with each dataset combining replicated Care and NoCare samples reared in otherwise identical conditions. Together, these resources position *N. vespilloides* as a powerful system for studying parental care and molecular development.

## Results

### Loss of care exposes genetic variation associated with body size

In the burying beetle, *N. vespilloides*, both parents raise their young on a small vertebrate carcass which they prepare prior to hatching (pre-hatching care) and provide direct post-hatching care through food provisioning and brood defence [2,3]. To determine whether parental care can buffer the phenotypic expression of population standing genetic variation, we removed parents at three separate time points to recreate three levels of care: Full Care, Medium Care and No Care (**Figure 1a**, **Methods**). We then measured adult body size across multiple individuals across families within each treatment. As expected [15,16], we found that the loss of care affected body size, with offspring reared without care (No Care) were significantly smaller in body size than both Full and Medium Care (**Figure 1b, Tables S1**). Despite similar overall total phenotypic variance across care treatments, variance partitioning differed strongly between groups (**Tables S1-2; Figure S2**). Specifically, offspring reared with No Care showed the greatest variance between families in average body size, and the least variance within families compared to Full and Mid Care offspring (**Figure 1, Table S1**). Moreover, we calculated broad-sense heritability estimates for each treatment which indicated that the proportion of total phenotypic variation due to genetic differences was greatest in the No Care group (**Figure 1c**), which is further consistent with parental care suppressing the phenotypic expression of standing genetic variation even when provided at moderate levels, and its removal revealing it.

**Figure 1.**
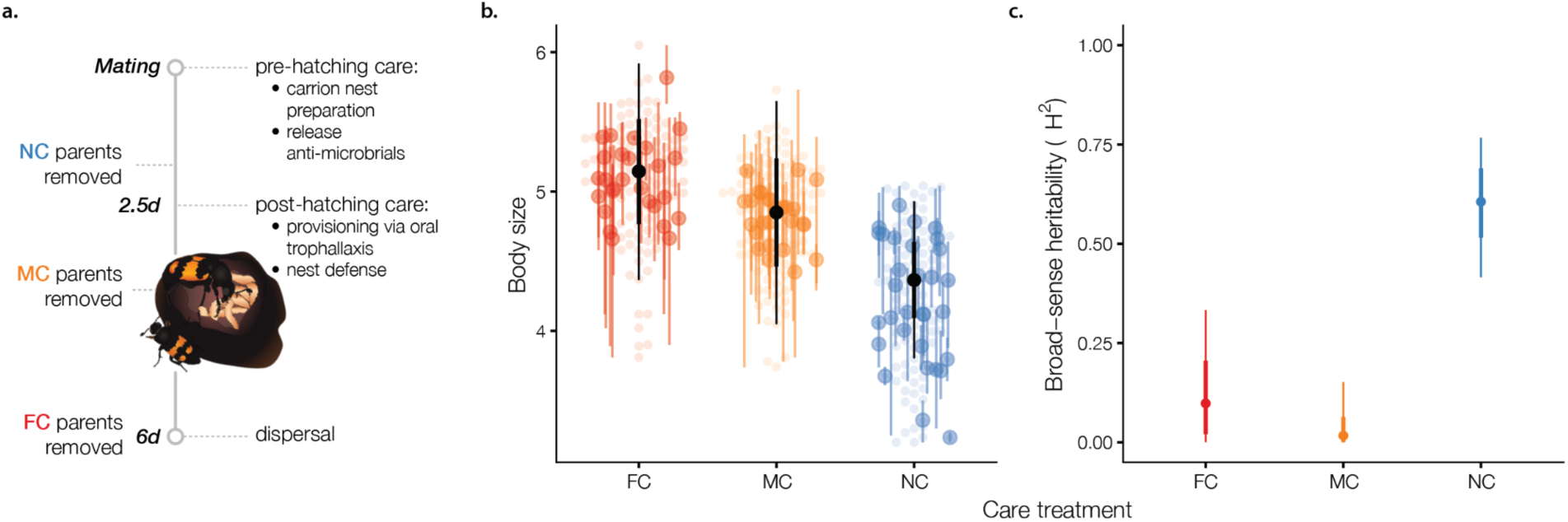
Parental care affects offspring body size. **a.** Schematic of the care manipulations. Parents were removed at three timepoints to generate No Care (NC; removed at 11h post-hatching), Medium Care (MC; removed at 24h post-hatching), and Full Care (FC; removed at larval dispersal) treatment groups, spanning pre- to post-hatching care. Figure adapted from [17]. **b.** Body size is lower on average in NC relative to MC and NC treatment groups. Black points and corresponding vertical lines show the median with 66% and 95% quantile intervals computed from draws of the posterior predictive distribution. Light coloured points show the raw data; coloured points with vertical lines show the raw means and ranges of body size for each family. **c.** Broad sense heritability (the ratio of among-family variance to the sum of among-family and residual variance) is greater in NC relative to MC and FC treatment groups. Points and vertical lines show the median with 66% and 95% quantile intervals from draws of the statistical model.

### A chromosome-scale genome assembly for *Nicrophorus vespilloides*

We sequenced and assembled the *Nicrophorus vespilloides* genome using high-coverage long reads together with proximity ligation data for scaffolding (**Methods**). The resulting assembly contains 7 chromosome-scale scaffolds spanning 202 Mb, as expected for this species [18], and a total of 14,340 genes of which 12,848 are protein-coding (98.4% complete BUSCO [19], **Figure S3, Tables S3-6**). Repeats account for 15.05% of the genome, with the majority arising from a recent burst of repeat expansion consisting mostly of LINE elements (**Figure S4**). We observe a strong collinearity with chromosomes of the sister species *Nicrophorus investigator* [18], further validating the quality of the assembly (**Figure S3**). Our chromosome-scale genomic resource comprehensively captures the *N. vespilloides* gene repertoire and associated regulatory domains, enabling rigorous analysis of molecular omics datasets.

### A single-nucleus atlas of the *N. vespilloide*s larval head

To elucidate the molecular mechanisms by which parental care might buffer development, we generated a single-nucleus atlas for head tissues from first instar larvae, comprising larvae from the two extreme care treatments: larvae reared with full parental care (Care) and larvae reared without (NoCare). After stringent filtering, the atlas comprises over 16,000 high-quality nuclei (**Figure 2a, Figure S5**), which we classified into 23 distinct cell types or states (**Methods**). We first broadly annotated cell-types based on well-established markers in *Drosophila* and other insects (**Figure 2b-c, Tables S7-S8**), uncovering a large fraction of epithelial cells (expressing grh, 56% of cells) likely involved in cuticle secretion (Hr38), muscle cells (Mhc, Mef2; 24%) and neuronal or glial cells (mature neurons: fne, Syt1, para; neuroglioblast/glia: pnt, repo, Gs2; 11%). We further detected rarer cell populations, including hemocytes (Hml and Ppn), gland cells (jhamt), tracheal cells (trh), fat body (apolpp, Apolptp), and an ECM-secretory population distinct from hemocytes and fat bodies, expressing specific collagens (CO1A2, COBA1).

**Figure 2:**
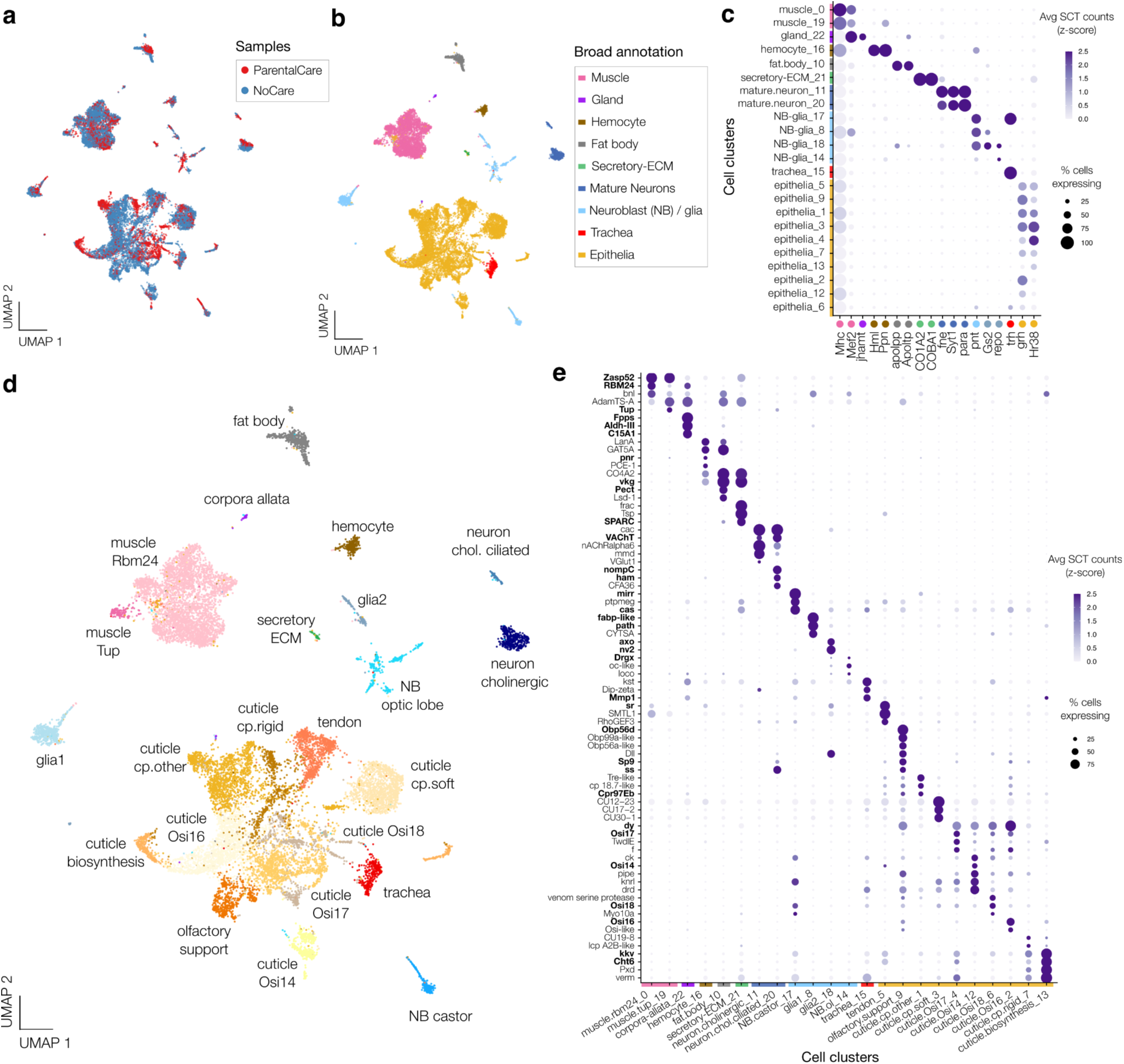
A single-nucleus atlas for the first instar larval head of *N. vespilloides*. **a.** Uniform manifold approximation and projection (UMAP) of integrated single-nucleus transcriptomes of larvae reared with (red) and without (blue) parental care. **b.** Broad annotation of cell populations visualised on the UMAP projection. **c.** Dot plot showing expression of selected broad markers supporting annotations presented in b. **d.** Specific annotation of the 23 cell populations visualised on the UMAP projection. **e.** Dot plot showing expression of selected specific marker genes underlying annotations presented in d. Key, previously experimentally validated markers are highlighted in bold (**Tables S7-S8**).

To more precisely annotate cells, we examined specific markers of our cell populations, focusing on previously experimentally validated genes (**Tables S7-S8, Figure 2d-e**). The two muscle populations express distinct markers, specifically the RBM24 beetle-specific muscle marker and the dorsal muscle marker Tup [20,21]. The ECM-secretory cells express SPARC and thus may correspond to the secretory cells of the lymph gland [22]. Co-expression of jhamt and juvenile hormone biosynthetic enzymes in gland cells (Fpps, Aldh-III, C15A1) confirms they are corpora allata cells. The two mature neuron populations appear predominantly cholinergic (VaCHt), one of which is associated with cilia genes and sensory functions (nompC, ham) [23,24]. Moreover, amongst other neuron-related cell populations, we identified two distinct glial population: glia1 (Gs2, path, fabp-like) and glia2 (Gs2, axo, nrv2) [25–29]; as well as two putative neuroglioblast lineages: NB castor (cas, mirr) and NB optic lobe (Drgx) [30–32]. Within epithelial cells, we identified the tendon cells mediating muscle-cuticle attachment expressing stripe (sr), as well as the olfactory support cells of the antennae expressing general odorant binding proteins and the antennae specification gene spineless (ss). The 8 remaining epithelial populations were primarily associated with cuticle formation and architecture. Notably, we identified chitin-synthesising cells, which we termed ‘cuticle biosynthesis’, and which expressed the chitin synthase (kkv) and associated chitin-modification enzyme (Cht6). Further, 3 cell populations express specific architectural chitin-binding cuticular proteins (cp), specifically soft cuticle proteins ‘cp.soft’, rigid ‘cp.rigid’ and unclassified ‘cp.other’ (**Figure S5, Table S9**) [33]. Lastly, the 4 remaining cell clusters all express dusky (dy) together with specific combinations of Osiris (Osi) genes, suggesting a role in nanopatterning of the cuticle and formation of specialized structures including pores, spinules or bristles [34,35].

These annotations were largely supported by direct comparisons with reference single-cell from the Fly Cell Atlas using Pesci [36] (**Figure S5**). Altogether, our atlas represents an unprecedented, curated resource to study beetle development, and forms a stepping stone to investigate molecular consequences associated with the loss of parental care.

### Hsp83 is up-regulated in larvae that received parental care

To investigate how the loss of parental care affects gene expression in *N. vespilloides* larvae, we performed differential gene expression tests on bulk [37] and single-nucleus transcriptomes of first instar larval head. We retained differentially expressed genes detected in both bulk and single-nucleus data, ensuring that our results reflect genuine expression shifts rather than cell composition or methodological artefacts (**Methods**). With this approach, we identified a total of 85 genes significantly up-regulated in Care samples and 172 genes up-regulated in No Care samples (**Figure 3a-b, Figure S6, Table S10**). Strikingly, Hsp83 was among the genes up-regulated in Care larvae, showing broad expression and significant up-regulation in over 10 cell clusters, including muscles, fat body, putative optic lobe neuroglioblasts and glial1 cells, and various epithelial populations (**Figure 3a-c**). Given Hsp83’s role in maintaining developmental protein homeostasis in the presence of genetic variation [12], this suggests that one mechanism through which parental care buffers genetic variation is by providing an environment that promotes Hsp83 expression. Conversely, the absence of care leads to reduced Hsp83 levels, disrupting protein homeostasis and resulting in a loss of buffering capacity. Supporting this, other broadly expressed genes upregulated in Care samples are also associated with homeostasis. These include notably the putative RNA chaperones Edc1-like and DHX30 [38,39], which regulate mRNA stability and translation.

**Figure 3:**
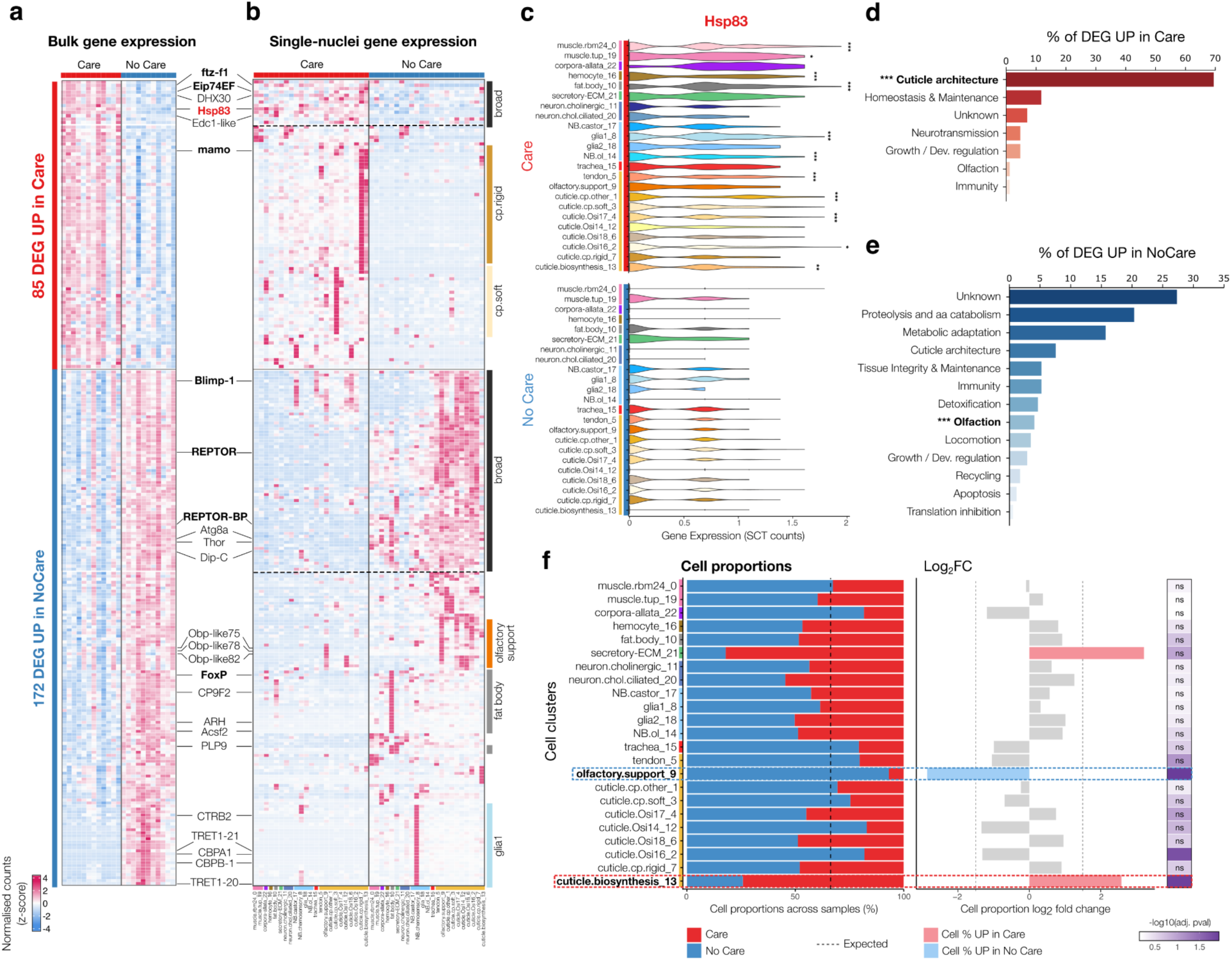
Changes in gene expression and cell type proportion driven by the loss of parental care. **a.** Expression of differentially expressed genes across bulk transcriptomics samples of 1^st^ instar larval heads (n=12 Care samples, n=11 NoCare samples). Counts were normalised for library size (counts per million) and transformed into z-score for visualisation. Key genes are highlighted on the right, with transcription factors in black bold. **b.** Expression of differentially expressed genes across cell clusters of the single-nucleus atlas, for Care (left) and NoCare (right) samples. Ribbons on the right highlight major expression patterns (broad or cluster-specific). Counts were SCT-normalised, adjusted for library size, logged and transformed into z-score. The colorbar is shared across (a) and (b) panels. **c.** Expression of Hsp83 across cell clusters, for Care (top) and NoCare (bottom) samples and shown as library adjusted logged SCT counts. Stars denote cell clusters in which Hsp83 is significantly up-regulated in Care (Wilcoxon BH-adjusted p-values, * p<0.05, ** p<0.01, ***p<0.001 and log_2_ fold-change>1). **d.** Functions of differentially expressed genes up-regulated in Care, with functions significantly enriched in Gene Ontology tests shown in bold (**Figure S6**). **e.** Functions of differentially expressed genes up-regulated in NoCare, as in (d). **f.** Differences in cell type proportions across Care and NoCare samples, with dotted boxes highlighting cells significantly more abundant in Care (red) or NoCare (blue) (BH-adjusted bootstrap-based p-values [40], p <0.05 and log_2_ fold-change>1.5).

### Parental care enables cuticle investment, whereas its loss induces a starvation stress response

To explore the functions of differentially expressed genes beyond individual candidates, we combined gene list curation and Gene Ontology enrichment approaches. We found that cuticle-related functions were significantly over-represented among Care up-regulated genes, accounting for 70% of up-regulated genes (**Figure 3d, S6**). Specifically, these included soft and rigid architectural chitin-binding proteins, which were expressed in corresponding soft and rigid cuticle-secreting cell populations. Consistently, the only up-regulated transcription factors identified were ftz-f1, Eip74EF and mamo, which are implicated in cuticular development and ecdysone-mediated growth [41–43].

Conversely, the most abundant functions among NoCare up-regulated genes were primarily associated with starvation stress, including proteolysis, metabolic adaptation and olfaction, with olfaction detected as significantly over-represented (**Figures 3b, e, S6**). In line with these observations, the only transcription factors up-regulated across multiple cell clusters were REPTOR, REPTOR-BP, and Blimp-1. Together, REPTOR and REPTOR-BP mediate most transcriptional shifts following starvation-induced TOR inhibition, while Blimp-1 represses ftz-f1 to delay energy-intensive developmental processes [44,45]. Other broadly up-regulated genes include key mediators of metabolic adaptations under nutrient deprivation, such as Thor, Dip-C, and Atg8a, which maintain energy homeostasis through translation inhibition, protein catabolism and autophagic recycling of cellular components [46–48]. In addition, the starvation response was also reflected in cell-type specific gene expression. The fat body, olfactory support and putative glial1 cells exhibited the highest number of cell-specific up-regulated genes. In the fat body, these genes were associated with lipid mobilization (FoxP, ARH, Acsf2, PLP9) and detoxification (CP9F2). Olfactory support cells showed an up-regulation of several general odorant-binding protein genes (Obp-like). In putative glial1 cells, up-regulated genes revealed metabolic and protective adaptations to sustain neuronal survival, including trehalose uptake (TRET1) [49], immune protection (PGRP-LF/SB1) and activation of various proteases (Chy/CTR1, CBPB, CBPA1), which may facilitate amino acid recycling or neuromodulation to preserve brain function under stress.

Lastly, we examined how care status shapes cell-type composition. We detected two specific cell populations with significant abundance shifts: olfactory support cells were enriched in the NoCare group, whereas cuticle-synthesizing cells were more abundant in the Care group (**Methods**, **Figure 3f**). Together with the up-regulation of Obp-like genes reported above, the increase in olfactory support cells likely represent an adaptation for locating food in the absence of parents [50]. Similarly, the reduction in cuticle-synthesizing cells in NoCare may reflect a lack of sufficient energy to fuel this metabolically expensive process.

Collectively, these findings suggest food provisioning is a critical component of parental care during early *N. vespilloides* development, sustaining overall growth and specifically driving continuous cuticle formation. Despite identical carcass access, the direct provision of pre-digested food by parents likely confers a crucial advantage.

### Distinct contributions of promoters and enhancers

To elucidate the regulatory changes driving differential gene expression, we performed genome-wide profiling of chromatin accessibility (ATAC-seq) and H3K4me3 histone mark enrichment (CUT&Tag) in head tissues of first-instar larvae, using four replicates for each care condition (**Figure S7, Table S11**). Following ATAC-seq peak calling, we defined a consensus set of 27,149 putative cis-regulatory elements which we classified into 21,139 putative enhancers and 6,010 promoters through overlaps with H3K4me3 peaks [51] (**Figure 4a, Table S12**). We confirmed that both classes displayed expected distances relative to their nearest TSS (**Figure 4b**).

**Figure 4:**
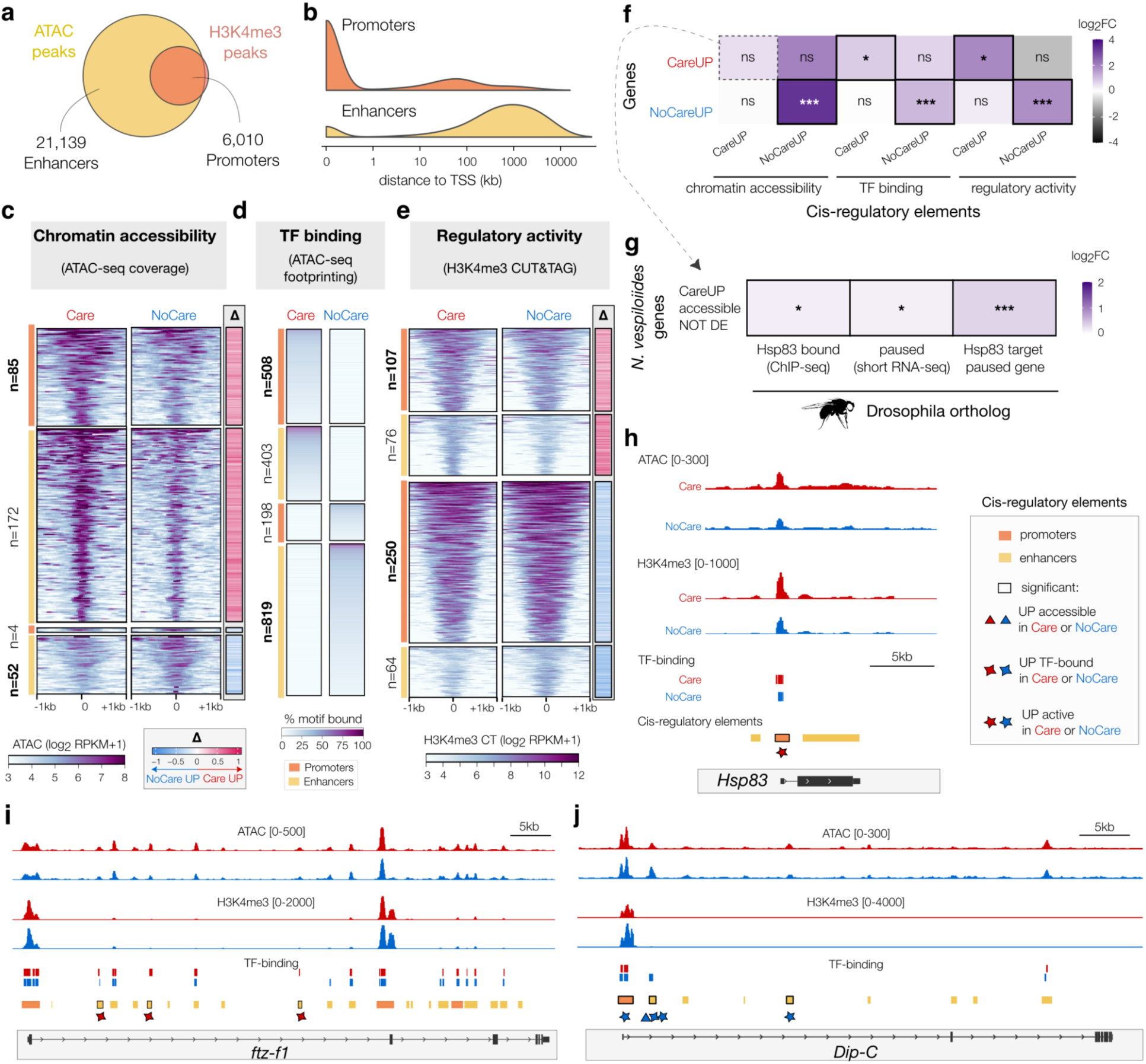
Regulatory changes driving gene expression shifts following the loss of parental care. **a.** Number of promoters and enhancers defined using chromatin accessibility and H3K4me3 histone mark enrichment. **b.** Distance of promoters and enhancers to their nearest transcription start site (TSS), confirming that most promoters are within 1 kb of their nearest TSS. **c**. Promoters and enhancers with significantly different accessibility across Care and NoCare groups. For visualisation, ATAC-seq read coverage (RPKM) was averaged across replicates for each group and log_2_-transformed. A normalised delta track is displayed on the right, showing relative differences in coverage between groups, specifically (Care-NoCare) / max(Care, NoCare). **d.** Promoters and enhancers with differential TF binding across Care and NoCare groups, as predicted using footprinting [52] (**Methods**). **e.** Promoters and enhancers with significantly different accessibility across Care and NoCare groups, as in (c), but showing H3K4me3 CUT & Tag read coverage. **f.** Association tests between differentially expressed genes and differential cis-regulatory elements (GREAT test hypergeometric p-values * p<0.05, ** p<0.01, ***p<0.001). **g.** Association tests between *N. vespilloides* genes with higher chromatin accessibility in Care without differential expression and *D. melanogaster* Hsp83 target and paused genes, using 1-1 orthologs (hypergeometric p-values * p<0.05, ** p<0.01, ***p<0.001; **Methods**). **h**. Coverage tracks around the Hsp83 CareUP gene. Coverage tracks are shown as RPKM and scaled to the tracks with highest magnitude. In the cis-regulatory element track, promoters (orange) and enhancers (yellow) with differential accessibility (triangle, red for CareUP, blue NoCareUP), TF-binding (4-branches star) or activity (five-branches star) are highlighted by a black box. **i.** Coverage tracks around the CareUP ftz-f1 TF gene, as in (h). **j.** Coverage tracks around the NoCareUP Dip-C gene, as in (h).

We investigated changes in the genome-wide regulatory landscapes of Care and NoCare larvae, examining three properties of consensus regulatory elements: chromatin accessibility (measured via ATAC-seq signal), transcription factor binding (ATAC-seq footprinting) and regulatory activity (H3K4me3 histone mark enrichment) (**Methods, Figure 4c-e**). Globally, we detected only subtle changes in chromatin accessibility and regulatory activity across conditions, reflecting small changes in signal magnitude rather than the actual gain or loss of regulatory elements. These results were robust to alternative peak-calling and statistical testing methods (**Methods**, **Figure S8**). Despite this global stability, ATAC-seq footprinting using TOBIAS [52] revealed pronounced changes in transcription factor (TF) occupancy between Care and NoCare larvae: mirroring biological functions observed in our gene expression data, top-ranked TFs enriched in NoCare larvae regulate nutrient-stress responses and neuronal lineage protection (e.g. l(3)neo38, GATA, klu, SREBP) [53–56], whereas those in Care larvae drive general developmental processes (e.g. bab1, fru, vis) [57–59] (**Table S13**).

Interestingly, promoters and enhancers contributed distinctly to Care and NoCare regulatory landscapes. Indeed, although promoters constitute a minority of all consensus regulatory elements (22%), they accounted for a disproportionate share of regions with increased accessibility (33%) and higher TF binding in Care larvae (Fisher’s exact test, both p < 0.001). Conversely, enhancers predominated in NoCare larvae, accounting for the vast majority of elements showing increased accessibility (93%) and TF binding (81%) (p <0.05). These findings suggest a subtle shift in regulatory architecture: while larvae reared with parental care rely on constitutive promoter-driven regulation, losing care leads to the recruitment of an alternative, enhancer-driven stress response.

### Parental care promotes a more open chromatin state

To test whether distinct regulatory logics operate in Care and NoCare larvae, we correlated regulatory shifts with differential gene expression, assigning consensus regulatory elements to their target genes based on TSS proximity (**Methods**). We found that a large fraction (42%) of regulatory elements with increased chromatin accessibility in Care larvae were associated with tRNA gene clusters. Although not captured by our mRNA-seq data, tRNA expression might be concordantly higher in Care larvae, reflecting a downregulation in NoCare individuals as part of their starvation stress response [60]. Restricting our analysis to elements associated with protein-coding genes, we found that all categories of regulatory elements were significantly associated with their respective differentially expressed gene sets, with one notable exception: regulatory elements with higher accessibility in Care larvae were not significantly associated with differential Care UP protein-coding genes (**Figure 4f**). Given that Hsp83 interacts with RNA polymerase II to maintain transcriptional pausing [61], we tested whether pausing could explain why many genes exhibit more accessible chromatin in Care larvae without corresponding changes in expression. Using published *Drosophila* Hsp83 ChIP-seq and short-capped RNA-seq data from embryonic cells as proxies for potential developmental Hsp83 targets and paused genes [61,62], we found that the accessible, non-differentially expressed genes in Care larvae significantly overlap with these putative Hsp83-mediated paused loci (**Table S14, Figure 4g**).

These results hint at a potentially more flexible and loose regulatory landscape in Care as opposed to NoCare larvae. Reflecting this, we found that Care UP genes were less likely than NoCare UP genes to be associated with concordant regulatory shifts across all three epigenomic layers (**Figures 4h-j, S9**). For instance, whereas the Care UP genes Hsp83 and ftz-f1 display only significantly increased promoter activity and TF binding in enhancers, respectively (**Figure 4h-i**), the NoCare UP gene Dip-C is marked by increased regulatory activity, accessibility and TF binding (**Figure 4j**). Together, these results reveal distinct regulatory features of larvae reared with and without parental care. While the loss of parental care triggers a coordinated, stress-driven activation of enhancers, Care larvae maintain a globally more open chromatin landscape that uncouples from immediate expression changes. This finding underscores that chromatin accessibility does not always drive active transcription. Instead, the more permissive chromatin state under care likely reflects Hsp83-mediated stabilisation, buffering development by priming access to transcription factors and RNA polymerase [63].

### Parental care promotes a more connected and redundant developmental regulatory network

To investigate whether parental care buffers phenotypic variation at the gene regulatory network level, we combined chromatin accessibility (ATAC-seq) and gene expression (RNA-seq) data to reconstruct genome-scale regulatory interaction networks for Care and NoCare larvae [64] (**Methods**). These regulatory networks capture the predicted genome-wide landscape of transcription factor-to-target-gene interactions in each care condition. Both the Care and NoCare networks displayed globally similar statistics (**Table S15**), sharing broadly the same regulatory factors. Moreover, as expected, condition-specific target genes corresponded to differentially expressed genes in each respective care condition (**Figure S10**).

Next, we identified the TFs most likely contributing to regulatory and expression differences between the Care and No Care conditions (Methods; Figure 5a). We identified these top TFs using the ANANSE influence score, which prioritises TFs that are strongly upregulated and/or control substantially more up-regulated target genes in one condition over the other (**Methods**). We then used these top 25 TFs to build a TF-TF subnetwork for each condition (**Figure 5b–f**). Consistent with the functions of the differentially expressed genes identified in our bulk and single-nucleus analysis (**Figure S10**), top Care TFs reflected an ecdysone-driven developmental growth programme, involving the ecdysone receptor and ecdysone-responsive TFs (EcR, ftz-f1, Eip74EF) as well as homeobox patterning TFs (btn, Sox14, vis, Dfd, slou) [42,65,66]. In contrast, top NoCare TFs were involved in metabolic stress response (foxo, foxP, fkh, SREBP) and possibly neuronal maintenance (vnd, onecut, pnt) [56,67,68]. Moreover, despite this method relying on motif accessibility rather than explicit motif footprinting, results were globally consistent with our footprinting analysis (**Table S13**).

**Figure 5.**
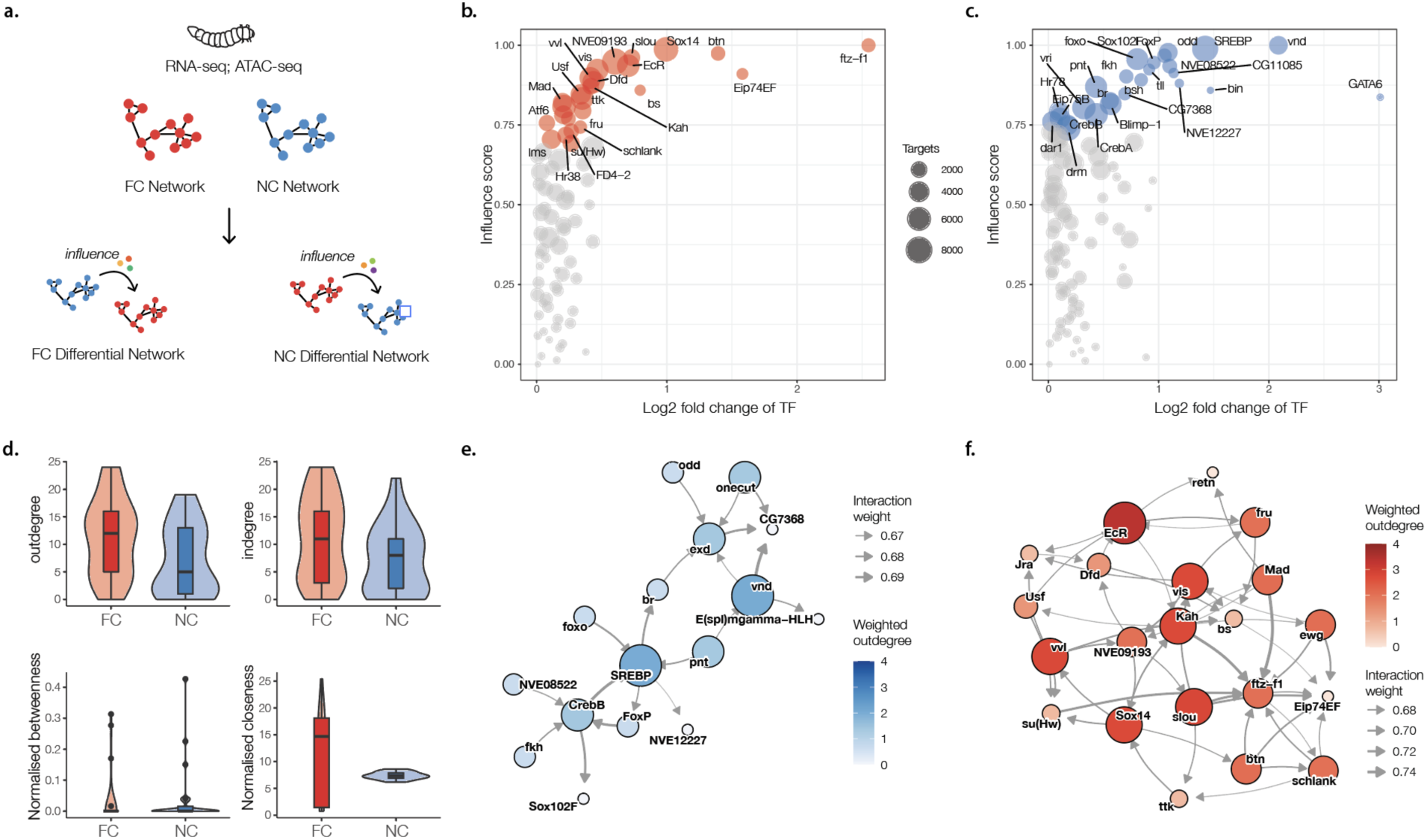
Regulatory networks under care. **a.** Schematic of the ANANSE analysis for networks of *cis*-regulatory interactions (adapted from [64]). Scatter plot of main transcription factors (TFs) driving the **b.** full care (FC) and **c.** no care (NC) networks. **d.** Violin and box plots comparing outdegree, indegree, normalised betweenness, and normalised closeness centrality of the top 30 influential TFs between FC and NC within the TF-TF subnetwork (Wilcoxon rank-sum tests; *p<0.05 and #p<0.10). **e.** Differential regulatory TF-TF subnetwork of the top influential TFs under FC, and **f.** NC. Node size and colour reflect weighted outdegree (sum of outgoing interaction scores to other top TFs); edge width reflects the predicted interaction strength of each individual TF-TF regulatory relationship.

We next analysed the structure of both condition-specific TF-TF subnetworks. The Care subnetwork showed significantly higher out-degree and in-degree, indicating greater mutual connectivity among its hub TFs (**Figure 5d**; out-degree: W = 403.5, p = 0.039; in-degree: W = 401.5, p = 0.043). Betweenness centrality, which measures how often a TF acts as a bottleneck, did not differ significantly between conditions (W = 254, p = 0.23). However, closeness centrality trended higher and showed substantially greater variance in the Care condition (W = 352, p = 0.054; **Figure 5d**). This pattern suggests that the Care network relies on a subset of globally connected coordinator TFs. Ultimately, this tighter, more distributed connectivity among Care hubs points to a more redundant regulatory architecture. This redundant network architecture highlights a potential mechanism through which parental care could act to buffer phenotypic development via gene regulatory interactions and their network structure.

## Discussion

Together, our findings demonstrate that parental care meets the criteria of a genetic capacitor. Classically, a genetic capacitor is defined as a molecular mechanism that masks the phenotypic expression of standing genetic variation, allowing it to accumulate, and releases it upon disruption [13,69]. Here, we show that a social behaviour, parental care, fulfils the same criteria: its presence suppresses among-family genetic variation in body size and reduces its heritability, whereas its loss unmasks this variation potentially exposing it to selection. These dynamics build directly upon our earlier finding that a history of care results in increased standing genetic variation. Moreover, at the molecular level, the buffering mechanisms we uncover align with classical models of genetic capacitance. We find that the canonical capacitor Hsp83 is upregulated under care and down-regulated upon its loss, alongside putative mRNA chaperones (Edc1-like, DHX30), implicating both proteostasis and post-transcriptional control as an important axis of developmental buffering.

Additionally, we identify distinct changes in the regulatory landscapes of larvae reared with or without parental care. Larvae reared with care maintain a globally more open, promoter-driven chromatin state, which is frequently uncoupled from immediate transcription and enriched for putative Hsp83-mediated paused loci. In contrast, the loss of care triggers a coordinated, enhancer-driven starvation–stress programme. At the network level, care establishes a redundant, densely connected architecture of transcription factor hubs. We propose that this network architecture may absorb among-family genetic variation to stabilise developmental phenotypes, which is disrupted under the stressful loss of care. This permissive, poised and redundant regulatory architecture under care is consistent with a model of genetic capacitance [12,13,69]. Taken together, care-mediated buffering therefore relies on the coordinated action of several regulatory layers, involving transcript and protein stabilisation, chromatin accessibility and gene network connectivity.

Interestingly, we found that specific cell types and tissues are differentially affected by the loss of parental care. For instance, our results suggest that larvae deprived of parental care mount a starvation response that redirects resources to prioritise neuronal maintenance and immediate survival at the expense of cuticle investment and growth. Thus, the loss of care may not expose genetic variation evenly across tissues, and variation underlying nervous system development may remain masked. Similarly, the broader effects of care loss are likely to vary across different developmental stages. Because our data capture only a single snapshot during the first instar, how these tissue trade-offs play out over time remains unknown. Care-deprived larvae do not differ markedly in overall developmental timing (time to dispersal or pupation time) [16], indicating that growth and developmental programmes might be globally reorganised instead of simply delayed. Tracking these dynamics across later developmental stages will be critical to determine whether larvae eventually recover from this initial starvation or maintain a continuous stress response throughout development and how this in turn affects development.

A central question is whether parental care and the variation it harbours could fuel adaptation to changing environments. At the genetic level, whether the variation released upon care loss represents a deleterious load or an adaptive reservoir depends on both population genetic architecture and environmental context. While adaptive examples of released cryptic variation remain rare in animals, notable examples have been documented in cavefish and red flour beetles [10,11]. For instance, disruption of Hsp90 (the vertebrate homolog of Hsp83) in surface-dwelling cavefish unmasks standing variation in eye size that, when selected, mirrors the reduced-eye phenotype evolved independently in cave-adapted populations [10]. Similarly, we previously showed that the loss of care can similarly unmask adaptive variants that are beneficial for adapting to a life without care [70]. Parental care could, therefore, act as a tunable buffer that conceals genetic variation under stable conditions and exposes it when environments change, thereby facilitating rapid adaptation. If care-mediated capacitance facilitates adaptation, it should ultimately translate into macroevolutionary patterns reflecting the evolutionary success of caring lineages. Consistent with this, parental care has repeatedly evolved across animal lineages, elaborate care/cooperation in birds is disproportionately associated with harsh and unpredictable environments [71], and caring fish lineages often display elevated diversification rates [72]. Ultimately, understanding these large-scale patterns requires uncovering their underlying mechanisms. By linking care-mediated buffering and release to specific molecular machinery across multiple scales, our work provides a mechanistic foundation to test how social behaviour shapes genome evolution, evolvability and population resilience to environmental change.

## Materials & Methods

### Beetle husbandry and the experimental manipulation of care

The beetles used in this experiment were derived from wild populations collected across six sites in Cambridgeshire, UK. Following a common garden generation, stock populations were established by interbreeding individuals collected across all six sites. Beetle pairs were bred by placing each pair in a separate breeding box (17 × 12 × 6 cm) with moist soil and a thawed mouse carcase (12-14g). We then placed each breeding box in a cupboard, and allowed parents to prepare the carcass and for the female to lay the clutch of eggs. To control for genetic background, individuals from each family were cross-paired with individuals from a genetically unrelated family, such that each sibling group was represented across multiple pairings (e.g., four males from Family 1 were each paired with a different female from Family 2). These pairs were distributed across three parental care treatments: 1) Full Care (FC), in which parents were left undisturbed; 2) Mid Care (MC), in which parents were removed 24 hours post-hatching; and 3) No Care (NC), in which parents were removed 53 hours post-pairing. At all life stages, individuals were kept at 21 °C. During breeding, pairs were kept in a constantly dark environment, but for all other life stages, they were kept on a 16L: 8D hour light cycle.

### Analysis of Body Size

At larval dispersal (8 days after pairing), we counted the number of dispersing larvae and weighed the entire brood. The dispersing larvae were placed in individual 2 × 2×2 cm cells within a plastic eclosion box (10 × 10 × 2 cm), covered with damp peat, and left to pupate. Newly eclosed adults were given a unique identifier, housed individually in plastic boxes (12 × 8 × 2 cm), and fed ground beef twice weekly until sexually mature (14 days post-eclosion), at which point pronotum width was measured as a proxy for body size [73]. We sampled up to 4 adults from each family (FC: n = 27 families, MC: n = 26, NC: n = 31) for a total sample size of 315 adults.

### Statistical analysis

We carried out the statistical analysis of body size in R v4.5.1 [74] using Bayesian models implemented in the package *brms* [75–77]. Additional R packages used for data handling, analysis and visualisation included tidyverse [78], DHARMa [79], tidybayes [80], ggdist [81], patchwork [82] and RColorBrewer [83].

Our model included fixed effects of care treatment, brood size (mean-centred), and their interaction. We also included covariates of sex and carcass weight (mean-centred and scaled to standard deviation units). Family ID was included as a random effect, with among-family variances estimated separately for each care treatment. It is important to note that among-family covariances were not estimated across care treatments as each family appeared in only one treatment group. We also estimated residual variances separately for each care treatment (again, without estimating covariances for the same reason). For variables of interest we compared the predictive ability of the model with and without that variable (or interaction) using leave-one-out (loo) cross-validation [52]. Although leave-one-out cross-validation did not identify the weak care treatment × brood size interaction (**Table S1; Figure S1**) as an important predictor (**Table S2**), we retained it in the final model because excluding it led to artificial inflation of among-family variance in the NC group, which would have confounded our primary inference.

We considered variation among families to represent genetic diversity within the population, and thus the proportion of total variation (after accounting for fixed effects) explained by among-family diversity in each treatment is an estimate of broad-sense heritability (i.e., H^2^ = V_G_/(V_G_+V_E_), where V_G_ represents among-family variance and V_E_ is the residual variance from the model). Models run via *brms* do not provide p-values but instead estimates with 95% credible intervals, which offer an indication of the effect size of variables in the model.

### Genome Assembly & Annotation

#### Genome assembly

High molecular weight DNA was extracted from a single female pupa using the MagAttract HMW DNA Kit (Qiagen) and we confirmed the presence of a 60kb peak using the Agilent 4200 TapeStation system. PacBio HiFi sequencing was performed by Novogene (Cambridge, UK) using the PacBio Revio platform, generating 52.9 Gb of HiFi reads (∼265× estimated coverage). The genome was assembled de novo using hifiasm v0.25.0-r726 [84] with default parameters optimized for HiFi reads. Haplotypic duplications were removed using purge_dups v1.2.6 (https://github.com/dfguan/purge_dups). Assembly quality and contamination were assessed using the nextflow DSL2 [85] BlobToolKit v0.9.9 pipeline [86] and contaminated contigs were removed prior to scaffolding and downstream analyses and annotation (**Figure S3**).

#### Scaffolding with HiC

Hi-C libraries were prepared using the Arima Genomics Hi-C kit using tissue from a single adult and sequenced on an Illumina NovaSeq 6000 platform to generate XX million 150-bp paired-end reads (PRJEB108640). Hi-C reads were processed using the Arima Genomics mapping pipeline (https://github.com/ArimaGenomics/mapping_pipeline). Briefly, reads were aligned to the draft assembly using bwa-mem2 v2.3 [87], filtered and used to scaffold the draft assembly with YaHS v1.2.2 [88]. MitoHiFi v3.2.3 [89,90] was used to identify and annotate the mitochondrial genome. Hi-C contact maps were generated using JuicerTools v1.22.01 [91] and visualized in Juicebox v2.15, which strongly supported the expected seven chromosomes (**Figure S3**) [92]. To evaluate assembly quality, we generated a K-mer spectrum plot using Merqury v1.3 [93] (**Figure S3**), which revealed a consensus quality (QV) score of 62.64 (i.e., fewer than one base-calling error per 1.5 million bases) and a K-mer completeness of 83.23%. The final assembly achieved a BUSCO completeness score of C:99.4% [S:97.9%, D:1.5%], F:0.1%, M:0.5% (n = 2,124, endopterygota_odb10) (**Table S4**).

#### Genome annotation

We used Liftoff v1.6.3 [94] to annotate the genome assembly by transferring gene models previously inferred by the NCBI EGAP annotation pipeline from the original *N. vespilloides* NCBI Refseq Assembly (GCF_001412225.1), which includes both protein-coding and non-coding genes. The gene annotation contained a total of 12,848 protein-coding genes, 149 pseudogenes, 351 lncRNA genes, 2 rRNA genes and 990 tRNA genes, and achieved a BUSCO completeness score of C:98.4% [S:97.1%, D:1.4%], F:0.7%, M:0.7% (n = 2,124, endopterygota_odb10) (**Table S4-5**). To facilitate functional annotation and naming of the predicted genes, we (i) identified one-to-one orthologs with *D. melanogaster* using reciprocal best BLAST hits, (ii) detected homologous proteins within the UniProt database via BLASTP searches, and (iii) identified functional domains using pfam_scan and InterProScan (**Table S6**). To facilitate functional interpretation of protein-coding genes, gene symbols were assigned using a hierarchical approach: *D. melanogaster* gene names were applied to genes with one-to-one orthologs (n = 7,075; 55%), UniProt names were assigned to the remaining genes with a valid BLASTP hit (n = 2,048; 16%) and the final unresolved fraction was designated with uncharacterized gene identifiers (n = 3,725; 29%). Putative transcription factors (TFs) were identified by screening the full predicted proteome and retaining all proteins containing an inferred sequence-specific DNA-binding [95] PFAM domain or InterPro entry [96,97]. To identify and classify genes encoding chitin-binding architectural cuticle proteins, we selected candidate proteins containing the ‘Chitin_bind_4’ PFAM domain (representing the arthropod cuticle Rebers and Riddiford [R&R] consensus) and subsequently classified them into soft (RR-1) and rigid (RR-2) cuticle proteins using CuticleDB [33] (**Table S9**). Comparison of the assembly and annotation statistics with previous *N. vespilloides* genome assemblies are reported in **Tables S3-5**.

#### Repeat annotation

We used EarlGrey v7.2.0 [98,99] to perform *de novo* annotation of transposable and repetitive elements. EarlGrey employs RepeatModeler2 [100] for *de novo* repeat family identification, followed by iterative consensus extension and refinement using the Blast, Extract, Align, Trim (BEAT) pipeline to generate full-length TE consensus sequences followed by a series of curation steps to defragment, remove overlapping, and filter spurious annotations, before producing a final genome annotation with RepeatMasker [101,102] (**Figure S4**).

#### Synteny analysis

We used the Eco-Flow nextflow [85] synteny pipeline (github.com/Eco-Flow/synteny) to perform pairwise macrosynteny comparison between the *N. investigator* (GCA_963457615.1) [18] and *N. vespilloides* genomes (**Figure S3**). Briefly, protein-coding sequences were extracted from genome annotations and pairwise orthologous anchor pairs were identified using jcvi [103], with syntenic blocks constructed using MCScanX [104].

### Bulk transcriptomic analyses

We reprocessed bulk transcriptomic libraries from our previous study (GSE171776) of larvae either receiving care or no care, where each library was constructed from four larval heads from a single family (11–12 libraries per group) [37]. Data was processed using nf-core/rnaseq v3.21.0 [105] from the nf-core collection of workflows [106], utilising reproducible software environments from the Bioconda [107] and Biocontainers [108] projects. The pipeline was executed with Nextflow v25.10.0 [85]. Briefly, reads were quality filtered using Trimgalore v0.6.10 [109] and mapped to the genome assembly using STAR v2.7.11b [110] in two-pass mode. Gene-level read counts were then quantified using Salmon v1.10.3 [111] in alignment-based mode. Alignment rates to the new genome assembly reported here averaged 96.96% (SD 0.79). We performed differential expression analyses using DESeq2 v1.44.0 [112] in R v4.4.0 [74] where we considered genes with an absolute log2FoldChange > 1 and an FDR adjusted p-value of < 0.05 to be differentially expressed.

### Single-nucleus transcriptomics (snRNA-seq)

For each library, larval heads from 2–3 families were pooled (10 heads total) to ensure a sufficient number of nuclei.

#### Nuclei isolation

Before starting the nuclei extractions, ultracentrifuge SW 41 Ti Swinging-Bucket Rotors (Beckman Coulter) were cooled down in the Optima XPN-90 Ultracentrifuge (Beckman Coulter) at 4 oC to prevent degradation of RNA. Tissue was homogenised using 2 ml Tissue Grinder Homogenisation Glass Vessel (VWR International) and 2 ml Tissue Grinder Plunger (VWR International) containing 2ml ice-cold HB (Homogenisation Buffer: 320mM Sucrose, 5mM Cacl2, 3mM Mg Acetate, 10mM Tris pH 7,4, 0.1mM EDTA, 0.1% NP40, 0.0%1 Digitonin, 0.1 mM PMSF (Sigma), 1mM β-mercaptoethanol). Tissue was homogenised with 250 gentle strokes. 2ml of the homogenate were transferred to an ice-cold falcon tube, with additional 650 µl HB added, followed by adding 2.65 ml Gradient Medium (GM: 5mM Cacl2, 50% Optiprep, 3mM MgCl2, 10mM Tris pH 7,4, 0.1 mM PMSF, 1mM β-mercaptoethanol). The mixture was gently mixed by flipping, without vortexing. Ultracentrifuge tubes (Beckman Coulter) were first layered with 4 ml of Cushion buffer (2.175ml Optiprep and 2325 of ODN buffer: 150mM KCl, 30mM MgCl2, 50mM Tris pH 7,4, 250mM Sucrose), followed by adding 5.25 ml of homogenised sample in HB buffer. The samples were balanced by weight and then spun in an ultracentrifuge at 7700 RPM at 4 oC for 30 minutes. After centrifugation, the supernatant was carefully removed without disturbing the nuclei pellet using a Pasteur pipette. The nuclei were then resuspended in RNaseq Wash Buffer (1.79mL PBS without Calcium and Magnesium (Invitrogen), 1% BSA (Miltenyi Biotec) and 0.2U/µl RNase In RNAse inhibitor (Promega). Resuspended nuclei were filtered through 5 ml Round Bottom Polystyrene Test Tubes with Cell Strainer (SLS). The sample was then centrifuged with the table-top centrifuge (Eppendorf) at 500 RCF, 4 oC for 5 minutes. The supernatant was removed and the nuclei were resuspended again and spun down with the same conditions four times. Finally, supernatant was removed and nuclei resuspended in 100 µl of RNaseq Wash Buffer in DNA LoBind microcentrifuge tubes (Eppendorf). Nuclei were counted using C-Chip Neubauer Hemocytometer (Cambridge Bioscience) and light microscopy (Evos XL Core, Thermofisher), as well as LUNA-FL cell counter (Logos Biosystems) with Propidium Iodide/Acridine Orange (Invitrogen) staining.

#### Single-nucleus library preparation and sequencing

All samples were processed as per Chromium Next GEM Single Cell 3ʹ Reagent Kits v3.1 (Dual Index). Following manufacturer’s guidelines, the samples were processed to target 10,000 nuclei per sample. We performed 11 cycles of cDNA amplification and 13 cycles of final indexing PCR. The concentrations and quality of cDNA and indexed final libraries were measured using D5000 ScreenTape Assay and D1000 ScreenTape Assay, respectively, on a TapeStation 4200 instrument (Agilent). Samples were sequenced on an Illumina NextSeq-2000 instrument using a P2 100-cycle flow cell and cartridge.

#### snRNA-seq read mapping and droplet filtering

Reads were mapped to the transcriptome using 10x Genomic CellRanger v9.0.1 [113], taking introns into account with the option --include-introns=true. We used QClus [114] to filter out low-quality droplets for each sample. Briefly, QClus leverages QC metrics specifically tailored to single-nucleus data (% of reads from unspliced transcripts, expression of nuclear marker genes) in addition to classical metrics (% of mitochondrial counts, total number of UMIs and expressed genes), to cluster and filter empty or highly-contaminated droplets without having to set *a priori* thresholds. QClus further uses the Scrublet algorithm [115] to identify and remove putative doublets. After initial dimensionality reduction and clustering in Seurat v5 [116], we further filtered out all nuclei from a cluster which was driven by low-quality nuclei (notably displaying low total number of UMIs). After these two highly-stringent filtering steps, our final dataset contains a total of 16,201 high-quality nuclei (10,742 nuclei from ‘NoCare’ larval heads and 5,459 from ‘Care’ larval heads) expressing 10,720 genes (**Figure S5**).

#### snRNA-seq data analysis

The final 16,201 nuclei dataset was analysed in Seurat v5 [116]. We first normalised counts within each sample using SCT transformation (v2), to account for cell-level differences in sequencing depth and stabilise variance across genes. To integrate datasets across samples, we used Seurat’s CCA approach. Briefly, we identified the top 3,000 shared variable genes on the basis of their Pearson’s residuals in the SCT models (i.e. with SelectIntegrationFeatures on the SCT assay, followed by PrepSCTIntegration), next, reducing datasets to these top variable features, we projected cells into a shared CCA space to identify and filter integration anchors (FindIntegrationAnchors), and ultimately aligned the two datasets (IntegrateData). We used Seurat’s Louvain algorithm to cluster nuclei. Specifically, we performed PCA on the integrated dataset, retained the top 100 Principal Components as suggested suitable from ElbowPlot inspection, constructed a nearest neighbor graph comprising all groups of 20 nearest cells based on euclidean distance in this reduced PCA space (FindNeighbors with dims1:100), and finally used the Louvain algorithm to identify clusters that optimize the modularity score. Our final resolution for clustering was set to 0.4, resulting in a total of 23 cell clusters. For visualisation purposes, we computed a UMAP from the top 100 Principals Components, using 30 neighbors.

We used a combination of approaches for the annotation of cell clusters: inspection of the expression of previously validated marker genes in drosophila or beetle (**Table S7**), direct single-cell gene expression comparison with relevant dataset from the Fly Cell Atlas (head, trachea, antenna and proboscis) [117] using Pesci v0.1.0 [36] (**Figure S5**) and literature search on cluster-specific marker genes (**Tables S7-S8**). To identify cluster-specific marker genes from our single-nucleus data, we first applied PrepSCTFindMarkers following sample merging to normalize SCT-corrected counts across Care and NoCare datasets. We then performed Wilcoxon Rank Sum tests using Seurat’s FindAllMarkers, retaining genes expressed in >5% of cells in either the foreground or background clusters with an average log_2_FoldChange > 1 and a Bonferroni-corrected p-value < 0.05. Similarly, we performed differential expression analyses across nuclei of ‘NoCare’ and ‘Care’ samples for each cluster using Wilcoxon Rank Sum tests on SCT corrected counts using the same parameters, but allowing for negative log2Foldchange, i.e. filtering for absolute(log_2_FoldChange) > 1. To mitigate single-cell DE statistical limitations [118], we only retained genes identified as differentially expressed in both the bulk and single-cell datasets. Unless otherwise specified, gene expression is presented using logged, library-size-normalized SCT counts in violin plot representations, and as the z-score of these averaged normalized counts in dotplot and heatmap representations.

We used scanpro v0.4.1 [40] to test for differences in cell type abundance across Care and No Care groups. Specifically, we used scanpro’s bootstrapping method with default parameters and applied the recommended arcsin square root transformation to stabilize the variance of cell proportions. We retained as differentially abundant all cell types with absolute(log_2_FoldChange) > 1.5 and adjusted p-value<0.05.

#### Gene Ontology enrichment tests

We used eggNOG-mapper [119] to automatically annotate *N. vespilloides* genes with Gene Ontology (GO) terms from the Biological Process, Molecular Function and Cellular Component domains. We used the enricher function from the ClusterProfiler v4.18.4 [120] R package to conduct hypergeometric enrichment tests on foreground sets of differentially expressed genes against the whole-genome background, applying a BH-adjusted p-value threshold of p < 0.05 (**Figure S6**).

### H3K4me3 histone mark enrichment using Cut&Tag assays

#### Library preparation

Larval heads were removed and bisected prior to homogenisation in 1X phosphate-buffered saline (PBS) on ice. Cell counts were determined before proceeding with CUT&Tag reactions, with 40,000–60,000 cells used per reaction. CUT&Tag was performed according to the manufacturer’s instructions using the CUT&Tag-IT Assay Kit (Active Motif). Assays were performed using an antibody targeting the histone H3K4me3 modification (cat no. 39160; Active Motif) with rabbit IgG serving as a negative control. Library quality and nucleosomal periodicity was assessed using an Agilent TapeStation prior to sequencing.

#### Sequencing and read mapping

Cut&Tag libraries were sequenced by Novogene (Cambridge, UK) on an Illumina NovaSeq X Plus platform, generating paired-end 150 bp reads to a target depth of ∼13 million reads per sample. Reads quality filtered and trimmed using TrimGalore v0.6.10 [109] and mapped to the genome assembly using the bowtie2 v2.3.5.1 [121] aligner using the parameters: local alignment mode (“--local --very-sensitive-local”), no mixed or discordant read pairs (--no-mixed --no-discordant), and a fragment size range of 10–700 bp (-I 10 -X 700) (**Figure S7, Table S11**). Resulting alignments were filtered for MAPQ ≥20 and coordinate-sorted using samtools [122].

#### Peak calling

To identify regions of H3K4me3 enrichment, peaks were called collectively across all samples using Genrich v0.6.1 (https://github.com/jsh58/Genrich/) To specifically enrich for robust promoter peaks and filter out low-magnitude signals characteristic of enhancers, we retained peaks with a stringent q-value threshold of q < 10^−18^, which effectively enriches for TSS-proximal peaks (**Figure 3**, see ***Definition of promoters and enhancers***). Although traditionally a promoter mark, low H3K4me3 signal is also captured at enhancers; we therefore leveraged H3K4me3 read counts across both element types to test for differential regulatory activity between the Care and NoCare groups (see ***Differential accessibility and differential activity tests***).

### Chromatin accessibility profiling using ATAC-seq

#### Library preparation

We performed assay for Transposase-Accessible Chromatin with sequencing (ATAC-seq) using a modified protocol based on Henning et al. [123]. Consistent with bulk and snRNA sampling, larvae from Care and NoCare groups were sampled 11h post-hatching for chromatin accessibility analysis. For each library, larval heads from 2–3 families were pooled (10 heads total) to ensure a sufficient number of nuclei.

Working stocks of double-stranded adapters (50 μM) were prepared by mixing equal volumes of Tn5ME-A and Tn5MErev adapters, and Tn5ME-B and Tn5MErev adapters separately. Adapters were annealed by incubation at 95°C for 5 min, followed by slow cooling to 65°C (0.1°C/sec) with a 5 min hold, then slow cooling to 4°C (0.1°C/sec). Annealed adapters were aliquoted and stored at −20°C until use. Prior to Tn5 loading, annealed adapters were thawed on ice and diluted to 35 μM with nuclease-free water. For each reaction, 0.5 μl of each adapter mix was combined with 1 μl Tn5 and 3 μl water in a 1.5 ml tube and incubated at 23°C for 60 min with shaking at 350 rpm.Nuclei were prepared concurrently. Ten larval heads per condition were dissected into 350 μl sucrose buffer (10 mM Tris-HCl, 250 mM D-sucrose, 1 mM MgCl₂, pH 7.5, supplemented with protease inhibitors) in a 1.5 ml tube and homogenised with 10 pestle strokes. The suspension was briefly incubated on ice to allow debris to sediment, and the supernatant was transferred to a fresh tube and centrifuged at 1,000 × *g* for 8 min at 4°C. The supernatant was discarded and the pellet resuspended in 100 μl lysis buffer (10 mM Tris-HCl, 10 mM NaCl, 3 mM MgCl₂, 0.1% IGEPAL CA-630, pH 7.5) and incubated on ice for 5 min. Nuclei were pelleted by centrifugation at 1,000 × *g* for 8 min at 4°C. For tagmentation, nuclei were resuspended directly in a 20 μl reaction containing 4× tagmentation buffer, dimethylformamide, nuclease-free water, and 5 μl adapter-loaded Tn5. The reaction was incubated at 55°C for 10 min and cooled to 10°C. Tagmentation was quenched by addition of 5 μl 0.2% SDS and incubation for 5 min at room temperature. DNA was purified using a Qiagen MinElute PCR Purification Kit prior to library amplification by PCR. Following amplification, libraries were purified again with a Qiagen MinElute kit and quantified by Qubit fluorometry and TapeStation analysis.

#### Sequencing and read mapping

ATAC-seq libraries were sequenced by Novogene (Cambridge, UK) on an Illumina NovaSeq X Plus platform, generating paired-end 150 bp reads to an average target depth of ∼50 million reads per sample. Reads quality filtered and trimmed using TrimGalore v0.6.10 [109] and mapped to our NVES_3.0 genome assembly using the bowtie2 v2.3.5.1 aligner [121] (**Figure S7, Table S11**).

#### Definition of promoters and enhancers

We used Genrich v0.6.1 (https://github.com/jsh58/Genrich/) to call consensus peaks using all ATAC-seq samples together, with a minimum mapping quality (MAPQ) of 13. This consensus peak set contained a total of 27,149 candidate regulatory elements, which we classified into promoters and enhancers based on overlap with H3K4me3 peaks. Specifically, we used bedtools intersect (v2.31.0) to intersect both sets of peaks and classified as promoter any ATAC-seq peak intersecting an H3K4me3 peak. To validate the robustness of of pooling all samples during the primary Genrich peak-calling step, we compared our results against two alternative pipelines, specifically: (i) calling peaks separately for each condition (Care or NoCare) with Genrich and (ii) calling peaks for each individual replicate with MACS2 [124] followed by merging of reproducible peaks from each condition into a consensus set (**Figure S8**).

#### Differential accessibility and differential activity tests

To perform differential accessibility analysis, we first counted ATAC-seq reads within consensus peaks using the multicov command in bedtools v2.31.0 [125] and used DESeq2 v1.44.0 [112] in R v4.4.0 [74] to identify differentially accessible peaks (using FDR corrected p<0.05). Alternative differential analyses methods as implemented in DiffBind v3.23 [126] recovered similar results (**Figure S8**). Similarly, for differential activity, we counted H3K4me3 reads within consensus peaks and performed differential analysis with DESeq2 using the same statistical threshold.

#### Regulatory element to gene associations and enrichment tests

We applied the default genomic association rule from GREAT [127] to link candidate regulatory elements to their putative target genes. Briefly, a proximal regulatory domain was defined for each gene (including both non-coding and protein-coding genes) by extending 5kb upstream and 1kb downstream of their Transcription Start Sites. These domains were then distally extended to the nearest proximal domain or up to a max distance of 1Mp. Candidate regulatory elements were then associated with any gene whose regulatory domain they intersected. To test for significant association between differentially expressed genes and differential peaks, we first restricted the consensus peak set to peaks associated with at least one protein-coding gene. We then implemented GREAT’s genomic hypergeometric test over genes using the total filtered consensus peak set as background.

#### Transcription factor footprinting analysis

To investigate genome-wide transcription factor (TF) binding dynamics, we performed footprinting analyses using TOBIAS v0.17.3 [128]. TOBIAS infers TF binding events by detecting localised DNA regions that are physically protected from transposase cleavage by bound proteins. To achieve the high sequencing depth required for robust footprinting, BAM files from biological replicates of each condition were merged and subsequently downsampled to an identical depth of N = 60 million reads. We used TOBIAS ATACorrect to correct per-base insertion signals for Tn5 transposase bias across each condition, using the *N. vespilloides* genome and the consensus ATAC peak set as inputs. We subsequently computed footprint scores across all peaks using TOBIAS ScoreBigwig with default parameters. Lastly, we used TOBIAS BINDetect with default parameters to predict binding events and identify TFs with significant differences in genomic occupancy across conditions. This analysis was performed using a custom-curated position weight matrix (PWM) motif file containing n=225 insect TF motifs. To generate this curated set, we filtered the insect non-redundant JASPAR 2026 database [129] to retain only motifs corresponding to *D. melanogaster* TFs with an ortholog in *N. vespilloides* itself predicted to encode a functional TF (see ***Genome Annotation***). BINDetect reports the proportion of bound and unbound sites for each TF motif in peaks, and for each TF tests for significant differences in binding scores between conditions. We retained TFs with an absolute Log_2_ fold-change > 0.1 and p-value < 10^−70^ (**Table S13**).

#### Differential TF binding

We identified ATAC-seq peaks with significant differences in transcription factor binding between the Care and NoCare conditions. To achieve this, we applied Fisher’s exact test to evaluate differences in the fraction of total motif base pairs predicted as bound via footprinting between conditions, for each ATAC-seq peak. Differentially bound peaks were retained using a strict threshold of absolute Log_2_ fold-change > 2 and BH-adjusted p-value < 0.01 (**Figure 3**, **Table S12**).

#### Analysis of gene regulatory interactions

We used ANANSE v0.5.1 [64] to construct genome-wide TF-gene regulatory networks for each condition (Care and NoCare). We first generated a custom TF gene–motif association database for *N. vespilloides* in a consistent manner with the custom motif database used for our footprinting analysis (see ***Transcription factor footprinting analysis***). To achieve this, we first used GimmeMotifs motif2factors [130] using the proteomes and TF annotations of *Drosophila melanogaster* (BDGP6.54) alongside several other invertebrate proteomes including *Tribolium castaneum (*Tcas5.2)*, Apis mellifera (*Amel_HAv3.1), *Anopheles gambiae* (AgamP4), *Nasonia vitripennis (*Nvit_2.0)*, Monomorium pharaonis (*ASM1337386v2) *and Ooceraea biroi (*Obir_v5.4). We subsequently filtered these predicted *N. vespilloides* TF gene–motif association to retain only high-confidence pairs where: (i) the *N. vespilloides* gene was predicted to encode a functional TF (see *Genome annotation*), (ii) the associated motif belonged to the insect non-redundant JASPAR 2026 motif database [129] and (iii) the *N. vespilloides* gene was orthologous to the *Drosophila* gene linked to that reference motif. This resulted in a total of n = 245 high-confidence TF gene-motif associations.

For each condition (Care and NoCare), we ran ANANSE binding to calculate genome-wide transcription factor binding probabilities across our consensus ATAC-seq peak set using the ATAC-seq bam alignment files, the consensus peak coordinates and our custom TF gene-motif associations as inputs. We then ran ANANSE network to create a network of TFs and their target genes (TGs) using condition-specific binding profiles and RNA-seq expression counts. The resulting network was imported into R [74] and analysed using iGraph [131]. To generate network statistics, we subsetted the network to retrieve edges of the top 5% (ANANSE score value, ‘prob’, above quantile 0.95) and subsequently computed network size, number of connections per target gene (TG) and per TF class centrality using modified custom wrappers for iGraph functions in R [132]. Lastly, we used ANANSE influence to calculate network rewiring between the Care and NoCare conditions by integrating the RNA-seq differential expression statistics generated by DESeq2 (see above). This analysis identified the most influential transcription factor nodes in each network, that is, transcription factors most likely to drive changes in expression across conditions. The resulting differential network topologies were visualised using a custom script built upon the R packages iGraph [131], tidyverse [78], ggraph [133] and scales [134].

### Analysis of *D. melanogaster* Hsp83 ChIP-seq data

Raw sequencing reads for Hsp83 ChIP-seq (GSE31226generated from untreated S2 cells derived from embryonic hemocytes [61] were downloaded from the NCBI Sequence Read Archive (SRA) using the SRA Toolkit v3.2.1 (https://github.com/ncbi/sra-tools). Adapter sequences were trimmed and quality-filtered using Trim Galore v0.6.10 [109] and reads were aligned to the *Drosophila melanogaster* (dm6) reference genome using Bowtie2 v2.2.5 [121] with parameters “--very-sensitive -k 1 --no-unal”, retaining only uniquely mapping reads with MAPQ ≥ 20. Following PCR duplicate removal using Picard v3.4.0 (http://broadinstitute.github.io/picard/), peaks were called independently for each replicate using MACS2 v2.2.9 [124]. Reproducible peaks were identified as those present in both replicates using bedtools intersect v2.31.0 [125]. Peaks were annotated to the nearest gene promoter or transcription start site (TSS) using HOMER v5.1 [135]. We filtered this resulting list of Hsp83-bound *D. melanogaster* genes to retain only genes with a 1-to-1 gene in *N. vespilloides* (see ***Genome Annotation***). Finally, we used hypergeometric tests to test for significant overlap between these putative Hsp83 target genes and genes associated with differentially accessible peaks in the Care or NoCare conditions (**Figures 3, S9**).

### Analysis of *D. melanogaster* short-capped RNA-seq data and polymerase pausing

Raw sequencing reads for short-capped RNA-seq datasets (3’ and 5’ end libraries, generated from untreated S2 cells derived from embryonic hemocytes) from Nechaev et al. 2010 (GSE18643) were downloaded using the SRA Toolkit v3.2.1. Short RNA reads were hard-trimmed to 26 nt from the 5’ end to match the original study’s processing pipeline using TrimGalore v0.6.10 [109], then aligned to dm6 using Bowtie2 v2.2.5 [121] with parameters “--very-sensitive -k 1 --no-unal”, retaining only uniquely mapping reads with MAPQ ≥ 20. Biological replicates were merged using samtools merge v1.11 [122] prior to downstream analysis. Read counts in transcription start site (TSS) windows were quantified using the bedtools multicov function [125]. A gene was classified as being paused by Pol II if it had at least one short RNA read (from either the 3’ or 5’ library) within 150 bp of its annotated TSS, as described previously [136]. Statistical tests for association between putative paused genes and genes associated with differentially accessible peaks were performed as described for Hsp83 putative targets (see ***Analysis of D. melanogaster Hsp83 ChIP-seq data***)

## Acknowledgements

We thank the Imbeault lab and Dr. Marco Geymonat for synthesising the Tn5 transposase and for generously providing sequencing adapters. This work was supported by a BBSRC Future Leaders Fellowship (BB/R01115X/1) and a Royal Society Small Research Grant (RGS\R1\191162) to RM. EP is supported by a Leverhulme Research grant (RPG-2025-274). FM is supported by a Royal Society University Research Fellowship (URF\R\241014). Work by EP and FM was also supported by the BBSRC grant BB/V01109X/1.

## Author Contributions

RM conceived and directed the research. RM, SJS, AT, DG, FM and MB-H conducted the experimental work. RMK provided access to beetle colonies. RM, TH and EP performed statistical and bioinformatic analyses. RM and EP led the writing of the paper. All authors edited and commented on the manuscript.

## References

1. Royle NJ, Smiseth PT, Kölliker M, editors. 2012 The evolution of parental care. First edition. Oxford, United Kingdom: Oxford University Press.

2. Potticary AL et al. 2024 Revisiting the ecology and evolution of burying beetle behavior (Staphylinidae: Silphinae). Ecology and Evolution 14, e70175. (doi:10.1002/ece3.70175)

3. Scott MP. 1998 THE ECOLOGY AND BEHAVIOR OF BURYING BEETLES. Annu. Rev. Entomol. 43, 595–618. (doi:10.1146/annurev.ento.43.1.595)

4. Snell-Rood EC, Burger M, Hutton Q, Moczek AP. 2016 Effects of parental care on the accumulation and release of cryptic genetic variation: review of mechanisms and a case study of dung beetles. Evol Ecol 30, 251–265. (doi:10.1007/s10682-015-9813-4)

5. Patterson C, Pilakouta N. 2024 Effects of Parental Care on the Magnitude of Inbreeding Depression: A Meta-Analysis in Fishes. The American Naturalist 203, E50–E62. (doi:10.1086/728001)

6. Pascoal S, Shimadzu H, Mashoodh R, Kilner RM. 2023 Parental care results in a greater mutation load, for which it is also a phenotypic antidote. Proc. R. Soc. B. 290, 20230115. (doi:10.1098/rspb.2023.0115)

7. Pilakouta N, Jamieson S, Moorad JA, Smiseth PT. 2015 Parental care buffers against inbreeding depression in burying beetles. Proc. Natl. Acad. Sci. U.S.A. 112, 8031–8035. (doi:10.1073/pnas.1500658112)

8. Queitsch C, Sangster TA, Lindquist S. 2002 Hsp90 as a capacitor of phenotypic variation. Nature 417, 618–624. (doi:10.1038/nature749)

9. Rutherford SL, Lindquist S. 1998 Hsp90 as a capacitor for morphological evolution. Nature 396, 336–342. (doi:10.1038/24550)

10. Rohner N, Jarosz DF, Kowalko JE, Yoshizawa M, Jeffery WR, Borowsky RL, Lindquist S, Tabin CJ. 2013 Cryptic Variation in Morphological Evolution: HSP90 as a Capacitor for Loss of Eyes in Cavefish. Science 342, 1372–1375. (doi:10.1126/science.1240276)

11. Sayed R, Şahin Ö, Errbii M, R R, Peuß R, Prüser T, Schrader L, Schulz NKE, Kurtz J. 2025 HSP90 as an evolutionary capacitor drives adaptive eye size reduction via atonal. Nat Commun 16, 9277. (doi:10.1038/s41467-025-65027-0)

12. Taipale M, Jarosz DF, Lindquist S. 2010 HSP90 at the hub of protein homeostasis: emerging mechanistic insights. Nat Rev Mol Cell Biol 11, 515–528. (doi:10.1038/nrm2918)

13. Aguilar-Rodríguez J, Jakobson CM, Jarosz DF. 2024 The Hsp90 Molecular Chaperone as a Global Modifier of the Genotype-Phenotype-Fitness Map: An Evolutionary Perspective. Journal of Molecular Biology 436, 168846. (doi:10.1016/j.jmb.2024.168846)

14. Levy SF, Siegal ML. 2008 Network Hubs Buffer Environmental Variation in Saccharomyces cerevisiae. PLoS Biol 6, e264. (doi:10.1371/journal.pbio.0060264)

15. Schrader M, Jarrett BJM, Kilner RM. 2022 Larval environmental conditions influence plasticity in resource use by adults in the burying beetle, *Nicrophorus vespilloides*. Evolution 76, 667–674. (doi:10.1111/evo.14339)

16. Eggert A-K, Reinking M, Müller JK. 1998 Parental care improves offspring survival and growth in burying beetles. Animal Behaviour 55, 97–107. (doi:10.1006/anbe.1997.0588)

17. Körner M, Steiger S, Shukla SP. 2023 Microbial management as a driver of parental care and family aggregations in carrion feeding insects. Front. Ecol. Evol. 11, 1252876. (doi:10.3389/fevo.2023.1252876)

18. Crowley LM et al. 2024 The genome sequence of the Banded Burying beetle, Nicrophorus investigator Zetterstedt, 1824. Wellcome Open Res 9, 343. (doi:10.12688/wellcomeopenres.21496.1)

19. Simão FA, Waterhouse RM, Ioannidis P, Kriventseva EV, Zdobnov EM. 2015 BUSCO: assessing genome assembly and annotation completeness with single-copy orthologs. Bioinformatics 31, 3210–3212. (doi:10.1093/bioinformatics/btv351)

20. Boukhatmi H, Frendo JL, Enriquez J, Crozatier M, Dubois L, Vincent A. 2012 Tup/Islet1 integrates time and position to specify muscle identity in Drosophila. Development 139, 3572–3582. (doi:10.1242/dev.083410)

21. Schmitt-Engel C et al. 2015 The iBeetle large-scale RNAi screen reveals gene functions for insect development and physiology. Nat Commun 6, 7822. (doi:10.1038/ncomms8822)

22. Irving P, Ubeda J-M, Doucet D, Troxler L, Lagueux M, Zachary D, Hoffmann JA, Hetru C, Meister M. 2005 New insights into Drosophila larval haemocyte functions through genome-wide analysis. Cellular Microbiology 7, 335–350. (doi:10.1111/j.1462-5822.2004.00462.x)

23. Cheng LE, Song W, Looger LL, Jan LY, Jan YN. 2010 The role of the TRP channel NompC in Drosophila larval and adult locomotion. Neuron 67, 373–380. (doi:10.1016/j.neuron.2010.07.004)

24. Moore AW, Jan LY, Jan YN. 2002 hamlet, a Binary Genetic Switch Between Single- and Multiple-Dendrite Neuron Morphology. Science 297, 1355–1358. (doi:10.1126/science.1072387)

25. Dong Q, Alvarez-Ochoa E, Nguyen P-K, Orih P, Fahey-Lozano N, Kosakamoto H, Obata F, Alexandre C, Cheng LY. 2025 The blood–brain barrier regulates brain tumor growth through the SLC36 amino acid transporter Pathetic in Drosophila. PLOS Biology 23, e3003496. (doi:10.1371/journal.pbio.3003496)

26. Freeman MR, Delrow J, Kim J, Johnson E, Doe CQ. 2003 Unwrapping Glial Biology: Gcm Target Genes Regulating Glial Development, Diversification, and Function. Neuron 38, 567–580. (doi:10.1016/S0896-6273(03)00289-7)

27. Ahn HJ, Jeon S-H, Kim SH. 2014 Expression of a set of glial cell-specific markers in the Drosophila embryonic central nervous system. BMB Rep 47, 354–359. (doi:10.5483/BMBRep.2014.47.6.177)

28. Pogodalla N, Kranenburg H, Rey S, Rodrigues S, Cardona A, Klämbt C. 2021 Drosophila ßHeavy-Spectrin is required in polarized ensheathing glia that form a diffusion-barrier around the neuropil. Nat Commun 12, 6357. (doi:10.1038/s41467-021-26462-x)

29. Yuan LL, Ganetzky B. 1999 A glial-neuronal signaling pathway revealed by mutations in a neurexin-related protein. Science 283, 1343–1345. (doi:10.1126/science.283.5406.1343)

30. Dillon NR, Doe CQ. 2024 Castor is a temporal transcription factor that specifies early born central complex neuron identity. Development 151, dev204318. (doi:10.1242/dev.204318)

31. Karim MR, Moore AW. 2011 Convergent Local Identity and Topographic Projection of Sensory Neurons. J Neurosci 31, 17017–17027. (doi:10.1523/JNEUROSCI.2815-11.2011)

32. Gizler L, Schneider K, Steigleder S, Benmaamar S, Schneuwly S, Rass M. 2024 The pivotal role of Drgx in survival, wiring and identity of T4/T5 neurons., 2024.03.12.584653. (doi:10.1101/2024.03.12.584653)

33. Magkrioti CK, Spyropoulos IC, Iconomidou VA, Willis JH, Hamodrakas SJ. 2004 cuticleDB: a relational database of Arthropod cuticular proteins. BMC Bioinformatics 5, 138. (doi:10.1186/1471-2105-5-138)

34. Sun Z, Inagaki S, Miyoshi K, Saito K, Hayashi S. 2024 Osiris gene family defines the cuticle nanopatterns of Drosophila. Genetics 227, iyae065. (doi:10.1093/genetics/iyae065)

35. Roch F, Alonso CR, Akam M. 2003 Drosophila miniature and dusky encode ZP proteins required for cytoskeletal reorganisation during wing morphogenesis. J Cell Sci 116, 1199–1207. (doi:10.1242/jcs.00298)

36. Parey E, Piovani L, Marlétaz F. 2026 eparey/pesci: Pesci v0.1.0. (doi:10.5281/zenodo.20595563)

37. Sarkies P, Westoby J, Kilner RM, Mashoodh R. 2024 Gene body methylation evolves during the sustained loss of parental care in the burying beetle. Nature Communications 15, 6606.

38. Neef DW, Thiele DJ. 2009 Enhancer of decapping proteins 1 and 2 are important for translation during heat stress in Saccharomyces cerevisiae. Molecular Microbiology 73, 1032–1042. (doi:10.1111/j.1365-2958.2009.06827.x)

39. Gokhale A et al. 2021 Mitochondrial Proteostasis Requires Genes Encoded in a Neurodevelopmental Syndrome Locus. J. Neurosci. 41, 6596–6616. (doi:10.1523/JNEUROSCI.2197-20.2021)

40. Alayoubi Y, Bentsen M, Looso M. 2024 Scanpro is a tool for robust proportion analysis of single-cell resolution data. Sci Rep 14, 15581. (doi:10.1038/s41598-024-66381-7)

41. Suzuki T, Kawasaki H, Yu RT, Ueda H, Umesono K. 2001 Segmentation gene product Fushi tarazu is an LXXLL motif-dependent coactivator for orphan receptor FTZ-F1. Proceedings of the National Academy of Sciences 98, 12403–12408. (doi:10.1073/pnas.221552998)

42. Uyehara CM, McKay DJ. 2019 Direct and widespread role for the nuclear receptor EcR in mediating the response to ecdysone in Drosophila. Proceedings of the National Academy of Sciences 116, 9893–9902. (doi:10.1073/pnas.1900343116)

43. Lai Y-W, Miyares RL, Liu L-Y, Chu S-Y, Lee T, Yu H-H. 2022 Hormone-controlled changes in the differentiation state of post-mitotic neurons. Current Biology 32, 2341–2348.e3. (doi:10.1016/j.cub.2022.04.027)

44. Tiebe M, Lutz M, De La Garza A, Buechling T, Boutros M, Teleman AA. 2015 REPTOR and REPTOR-BP Regulate Organismal Metabolism and Transcription Downstream of TORC1. Developmental Cell 33, 272–284. (doi:10.1016/j.devcel.2015.03.013)

45. Agawa Y et al. 2007 Drosophila Blimp-1 Is a Transient Transcriptional Repressor That Controls Timing of the Ecdysone-Induced Developmental Pathway. Molecular and Cellular Biology 27, 8739–8747. (doi:10.1128/MCB.01304-07)

46. Teleman AA, Chen Y-W, Cohen SM. 2005 4E-BP functions as a metabolic brake used under stress conditions but not during normal growth. Genes Dev. 19, 1844–1848. (doi:10.1101/gad.341505)

47. Hiraizumi K, Hourani CL, Zambarano MC, Freeman JE, Mathes KD. 1992 Dipeptidase-C in Drosophila melanogaster: genetic, ontogenetic, and tissue-specific variation. Biochem Genet 30, 603–624. (doi:10.1007/BF02399810)

48. Jacomin A-C et al. 2020 Regulation of Expression of Autophagy Genes by Atg8a-Interacting Partners Sequoia, YL-1, and Sir2 in Drosophila. Cell Rep 31, 107695. (doi:10.1016/j.celrep.2020.107695)

49. Hertenstein H, McMullen E, Weiler A, Volkenhoff A, Becker HM, Schirmeier S. 2021 Starvation-induced regulation of carbohydrate transport at the blood–brain barrier is TGF-β-signaling dependent. eLife 10, e62503. (doi:10.7554/eLife.62503)

50. Kordaczuk J, Wojda I. In press. Insect olfactory proteins: A comprehensive review with a special emphasis on the role of odorant-binding proteins in insect immunity. Insect Science n/a. (doi:10.1111/1744-7917.70204)

51. Schuettengruber B, Ganapathi M, Leblanc B, Portoso M, Jaschek R, Tolhuis B, Lohuizen M van, Tanay A, Cavalli G. 2009 Functional Anatomy of Polycomb and Trithorax Chromatin Landscapes in Drosophila Embryos. PLOS Biology 7, e1000013. (doi:10.1371/journal.pbio.1000013)

52. Bentsen M et al. 2020 ATAC-seq footprinting unravels kinetics of transcription factor binding during zygotic genome activation. Nat Commun 11, 4267. (doi:10.1038/s41467-020-18035-1)

53. Reddington JP et al. 2020 Lineage-Resolved Enhancer and Promoter Usage during a Time Course of Embryogenesis. Developmental Cell 55, 648–664.e9. (doi:10.1016/j.devcel.2020.10.009)

54. Kokki K et al. 2021 Metabolic gene regulation by Drosophila GATA transcription factor Grain. PLoS Genet 17, e1009855. (doi:10.1371/journal.pgen.1009855)

55. Xiao Q, Komori H, Lee C-Y. 2012 klumpfuss distinguishes stem cells from progenitor cells during asymmetric neuroblast division. Development 139, 2670–2680. (doi:10.1242/dev.081687)

56. Ugrankar-Banerjee R, Tran S, Srivastava S, Bowerman J, Paul B, Zacharias LG, Mathews TP, DeBerardinis RJ, Henne WM. 2026 SREBP governs a triglyceride:glycogen metabolic switch in Drosophila. 2024.05.13.593915. (doi:10.1101/2024.05.13.593915)

57. Vernes SC. 2014 Genome wide identification of Fruitless targets suggests a role in upregulating genes important for neural circuit formation. Sci Rep 4, 4412. (doi:10.1038/srep04412)

58. Couderc J-L, Godt D, Zollman S, Chen J, Li M, Tiong S, Cramton SE, Sahut-Barnola I, Laski FA. 2002 The bric à brac locus consists of two paralogous genes encoding BTB/POZ domain proteins and acts as a homeotic and morphogenetic regulator of imaginal development in Drosophila. Development 129, 2419–2433. (doi:10.1242/dev.129.10.2419)

59. Peng J, Wang B-J, Svetec N, Zhao L. 2025 Gene regulatory networks and essential transcription factors for de novo originated genes. Nat Ecol Evol 9, 1487–1498. (doi:10.1038/s41559-025-02747-y)

60. Wei Y, Tsang CK, Zheng XFS. 2009 Mechanisms of regulation of RNA polymerase III-dependent transcription by TORC1. EMBO J 28, 2220–2230. (doi:10.1038/emboj.2009.179)

61. Sawarkar R, Sievers C, Paro R. 2012 Hsp90 Globally Targets Paused RNA Polymerase to Regulate Gene Expression in Response to Environmental Stimuli. Cell 149, 807–818. (doi:10.1016/j.cell.2012.02.061)

62. Nechaev S, Fargo DC, dos Santos G, Liu L, Gao Y, Adelman K. 2010 Global Analysis of Short RNAs Reveals Widespread Promoter-Proximal Stalling and Arrest of Pol II in Drosophila. Science 327, 335–338. (doi:10.1126/science.1181421)

63. Tariq M, Nussbaumer U, Chen Y, Beisel C, Paro R. 2009 Trithorax requires Hsp90 for maintenance of active chromatin at sites of gene expression. Proceedings of the National Academy of Sciences 106, 1157–1162. (doi:10.1073/pnas.0809669106)

64. Xu Q, Georgiou G, Frölich S, van der Sande M, Veenstra GJC, Zhou H, van Heeringen SJ. 2021 ANANSE: an enhancer network-based computational approach for predicting key transcription factors in cell fate determination. Nucleic Acids Res 49, 7966–7985. (doi:10.1093/nar/gkab598)

65. Kirilly D, Gu Y, Huang Y, Wu Z, Bashirullah A, Low BC, Kolodkin AL, Wang H, Yu F. 2009 A genetic pathway composed of Sox14 and Mical governs severing of dendrites during pruning. Nat Neurosci 12, 1497–1505. (doi:10.1038/nn.2415)

66. Knirr S, Azpiazu N, Frasch M. 1999 The role of the NK-homeobox gene slouch (S59) in somatic muscle patterning. Development 126, 4525–4535. (doi:10.1242/dev.126.20.4525)

67. Zhu S, Barshow S, Wildonger J, Jan LY, Jan Y-N. 2011 Ets transcription factor Pointed promotes the generation of intermediate neural progenitors in Drosophila larval brains. Proceedings of the National Academy of Sciences 108, 20615–20620. (doi:10.1073/pnas.1118595109)

68. Nguyen DN, Rohrbaugh M, Lai Z. 2000 The Drosophila homolog of Onecut homeodomain proteins is a neural-specific transcriptional activator with a potential role in regulating neural differentiation. Mech Dev 97, 57–72. (doi:10.1016/s0925-4773(00)00431-7)

69. Bergman A, Siegal ML. 2003 Evolutionary capacitance as a general feature of complex gene networks. Nature 424, 549–552. (doi:10.1038/nature01765)

70. Mashoodh R, Trowsdale AT, Manica A, Kilner RM. 2023 Parental care shapes the evolution of molecular genetic variation. Evolution Letters 7, 379–388. (doi:10.1093/evlett/qrad039)

71. Jetz W, Rubenstein DR. 2011 Environmental Uncertainty and the Global Biogeography of Cooperative Breeding in Birds. Current Biology 21, 72–78. (doi:10.1016/j.cub.2010.11.075)

72. Helmstetter AJ, Papadopulos AST, Igea J, Van Dooren TJM, Leroi AM, Savolainen V. 2016 Viviparity stimulates diversification in an order of fish. Nat Commun 7, 11271. (doi:10.1038/ncomms11271)

73. Jarrett BJM, Schrader M, Rebar D, Houslay TM, Kilner RM. 2017 Social interactions within the family enhance the capacity for evolutionary change. bioRxiv, 115014. (doi:10.1101/115014)

74. R Core Team. 2024 R: A Language and Environment for Statistical Computing.

75. Bürkner P-C. 2018 Advanced Bayesian Multilevel Modeling with the R Package brms. The R Journal 10, 395. (doi:10.32614/RJ-2018-017)

76. Bürkner P-C. 2017 **brms** : An *R* Package for Bayesian Multilevel Models Using *Stan*. J. Stat. Soft. 80. (doi:10.18637/jss.v080.i01)

77. Bürkner P-C. 2021 Bayesian Item Response Modeling in *R* with **brms** and *Stan*. J. Stat. Soft. 100. (doi:10.18637/jss.v100.i05)

78. Wickham H et al. 2019 Welcome to the Tidyverse. JOSS 4, 1686. (doi:10.21105/joss.01686)

79. Hartig F. 2016 DHARMa: Residual Diagnostics for Hierarchical (Multi-Level / Mixed) Regression Models. 0.4.7. (doi:10.32614/CRAN.package.DHARMa)

80. Matthew Kay. 2024 tidybayes: Tidy Data and Geoms for Bayesian Models. (doi:10.5281/ZENODO.1308151)

81. Matthew Kay. 2025 ggdist: Visualizations of distributions and uncertainty. (doi:10.5281/ZENODO.3879620)

82. Pedersen TL. 2019 patchwork: The Composer of Plots. 1.3.2. (doi:10.32614/CRAN.package.patchwork)

83. Neuwirth E. 2002 RColorBrewer: ColorBrewer Palettes, 1.1-3. (doi:10.32614/CRAN.package.RColorBrewer)

84. Cheng H, Asri M, Lucas J, Koren S, Li H. 2024 Scalable telomere-to-telomere assembly for diploid and polyploid genomes with double graph. Nat Methods 21, 967–970. (doi:10.1038/s41592-024-02269-8)

85. Di Tommaso P, Chatzou M, Floden EW, Barja PP, Palumbo E, Notredame C. 2017 Nextflow enables reproducible computational workflows. Nat Biotechnol 35, 316–319. (doi:10.1038/nbt.3820)

86. Muffato M et al. 2026 sanger-tol/blobtoolkit v0.10.1 - Onix (patch 1). (doi:10.5281/ZENODO.7949058)

87. Vasimuddin Md, Misra S, Li H, Aluru S. 2019 Efficient Architecture-Aware Acceleration of BWA-MEM for Multicore Systems. In 2019 IEEE International Parallel and Distributed Processing Symposium (IPDPS), pp. 314–324. Rio de Janeiro, Brazil: IEEE. (doi:10.1109/IPDPS.2019.00041)

88. Zhou C, McCarthy SA, Durbin R. 2023 YaHS: yet another Hi-C scaffolding tool. Bioinformatics 39, btac808. (doi:10.1093/bioinformatics/btac808)

89. Uliano-Silva M et al. 2023 MitoHiFi: a python pipeline for mitochondrial genome assembly from PacBio high fidelity reads. BMC Bioinformatics 24, 288. (doi:10.1186/s12859-023-05385-y)

90. Allio R, Schomaker-Bastos A, Romiguier J, Prosdocimi F, Nabholz B, Delsuc F. 2020 MitoFinder: Efficient automated large-scale extraction of mitogenomic data in target enrichment phylogenomics. Molecular Ecology Resources 20, 892–905. (doi:10.1111/1755-0998.13160)

91. Durand NC, Shamim MS, Machol I, Rao SSP, Huntley MH, Lander ES, Aiden EL. 2016 Juicer Provides a One-Click System for Analyzing Loop-Resolution Hi-C Experiments. Cell Systems 3, 95–98. (doi:10.1016/j.cels.2016.07.002)

92. Vorontsov NN, Yadav JS, Lyapunova EA, Korablev VP, Yanina IYu. 1984 Comparative karyology of seven species of staphylinoid beetles (Polyphaga: Coleoptera). Genetica 63, 153–159. (doi:10.1007/BF00605900)

93. Rhie A, Walenz BP, Koren S, Phillippy AM. 2020 Merqury: reference-free quality, completeness, and phasing assessment for genome assemblies. Genome Biol 21, 245. (doi:10.1186/s13059-020-02134-9)

94. Shumate A, Salzberg SL. 2021 Liftoff: accurate mapping of gene annotations. Bioinformatics 37, 1639–1643. (doi:10.1093/bioinformatics/btaa1016)

95. Blitz IL, Paraiso KD, Patrushev I, Chiu WTY, Cho KWY, Gilchrist MJ. 2017 A catalog of Xenopus tropicalis transcription factors and their regional expression in the early gastrula stage embryo. Dev Biol 426, 409–417. (doi:10.1016/j.ydbio.2016.07.002)

96. Mistry J et al. 2021 Pfam: The protein families database in 2021. Nucleic Acids Res 49, D412–D419. (doi:10.1093/nar/gkaa913)

97. Blum M et al. 2025 InterPro: the protein sequence classification resource in 2025. Nucleic Acids Res 53, D444–D456. (doi:10.1093/nar/gkae1082)

98. Baril T, Galbraith J, Hayward A. 2026 Earl Grey. (doi:10.5281/ZENODO.5654615)

99. Baril T, Galbraith J, Hayward A. 2024 Earl Grey: A Fully Automated User-Friendly Transposable Element Annotation and Analysis Pipeline. Molecular Biology and Evolution 41, msae068. (doi:10.1093/molbev/msae068)

100. Flynn JM, Hubley R, Goubert C, Rosen J, Clark AG, Feschotte C, Smit AF. 2020 RepeatModeler2 for automated genomic discovery of transposable element families. Proc. Natl. Acad. Sci. U.S.A. 117, 9451–9457. (doi:10.1073/pnas.1921046117)

101. Tarailo-Graovac M, Chen N. 2009 Using RepeatMasker to Identify Repetitive Elements in Genomic Sequences. CP in Bioinformatics 25. (doi:10.1002/0471250953.bi0410s25)

102. AFA Smit, R Hubley, P Green. 2013 RepeatMasker Open-4.0.

103. Tang H et al. 2024 JCVI: A versatile toolkit for comparative genomics analysis. iMeta 3, e211. (doi:10.1002/imt2.211)

104. Wang Y et al. 2012 MCScanX: a toolkit for detection and evolutionary analysis of gene synteny and collinearity. Nucleic Acids Research 40, e49–e49. (doi:10.1093/nar/gkr1293)

105. Harshil Patel et al. 2026 nf-core/rnaseq: nf-core/rnaseq v3.23.0 - Gallium Gecko. (doi:10.5281/ZENODO.1400710)

106. Ewels PA, Peltzer A, Fillinger S, Patel H, Alneberg J, Wilm A, Garcia MU, Di Tommaso P, Nahnsen S. 2020 The nf-core framework for community-curated bioinformatics pipelines. Nat Biotechnol 38, 276–278. (doi:10.1038/s41587-020-0439-x)

107. The Bioconda Team, Grüning B, Dale R, Sjödin A, Chapman BA, Rowe J, Tomkins-Tinch CH, Valieris R, Köster J. 2018 Bioconda: sustainable and comprehensive software distribution for the life sciences. Nat Methods 15, 475–476. (doi:10.1038/s41592-018-0046-7)

108. Da Veiga Leprevost F et al. 2017 BioContainers: an open-source and community-driven framework for software standardization. Bioinformatics 33, 2580–2582. (doi:10.1093/bioinformatics/btx192)

109. Krueger F, James F, Ewels P, Afyounian E, Weinstein M, Schuster-Boeckler B, Hulselmans G, Sclamons. 2023 FelixKrueger/TrimGalore: v0.6.10 - add default decompression path. (doi:10.5281/ZENODO.7598955)

110. Dobin A, Davis CA, Schlesinger F, Drenkow J, Zaleski C, Jha S, Batut P, Chaisson M, Gingeras TR. 2013 STAR: ultrafast universal RNA-seq aligner. Bioinformatics 29, 15–21. (doi:10.1093/bioinformatics/bts635)

111. Patro R, Duggal G, Love MI, Irizarry RA, Kingsford C. 2017 Salmon provides fast and bias-aware quantification of transcript expression. Nat Methods 14, 417–419. (doi:10.1038/nmeth.4197)

112. Love MI, Huber W, Anders S. 2014 Moderated estimation of fold change and dispersion for RNA-seq data with DESeq2. Genome Biol 15, 550. (doi:10.1186/s13059-014-0550-8)

113. Zheng GXY et al. 2017 Massively parallel digital transcriptional profiling of single cells. Nat Commun 8, 14049. (doi:10.1038/ncomms14049)

114. Schmauch E et al. 2025 QClus: a droplet filtering algorithm for enhanced snRNA-seq data quality in challenging samples. Nucleic Acids Res 53, gkae1145. (doi:10.1093/nar/gkae1145)

115. Wolock SL, Lopez R, Klein AM. 2019 Scrublet: Computational Identification of Cell Doublets in Single-Cell Transcriptomic Data. Cell Syst 8, 281–291.e9. (doi:10.1016/j.cels.2018.11.005)

116. Hao Y et al. 2024 Dictionary learning for integrative, multimodal and scalable single-cell analysis. Nat Biotechnol 42, 293–304. (doi:10.1038/s41587-023-01767-y)

117. Li H et al. 2022 Fly Cell Atlas: A single-nucleus transcriptomic atlas of the adult fruit fly. Science 375, eabk2432. (doi:10.1126/science.abk2432)

118. Squair JW et al. 2021 Confronting false discoveries in single-cell differential expression. Nat Commun 12, 5692. (doi:10.1038/s41467-021-25960-2)

119. Cantalapiedra CP, Hernández-Plaza A, Letunic I, Bork P, Huerta-Cepas J. 2021 eggNOG-mapper v2: Functional Annotation, Orthology Assignments, and Domain Prediction at the Metagenomic Scale. Mol Biol Evol 38, 5825–5829. (doi:10.1093/molbev/msab293)

120. Yu G, Wang L-G, Han Y, He Q-Y. 2012 clusterProfiler: an R Package for Comparing Biological Themes Among Gene Clusters. OMICS 16, 284–287. (doi:10.1089/omi.2011.0118)

121. Langmead B, Salzberg SL. 2012 Fast gapped-read alignment with Bowtie 2. Nat Methods 9, 357–359. (doi:10.1038/nmeth.1923)

122. Danecek P et al. 2021 Twelve years of SAMtools and BCFtools. GigaScience 10, giab008. (doi:10.1093/gigascience/giab008)

123. Hennig BP, Velten L, Racke I, Tu CS, Thoms M, Rybin V, Besir H, Remans K, Steinmetz LM. 2018 Large-Scale Low-Cost NGS Library Preparation Using a Robust Tn5 Purification and Tagmentation Protocol. G3 Genes|Genomes|Genetics 8, 79–89. (doi:10.1534/g3.117.300257)

124. Zhang Y et al. 2008 Model-based Analysis of ChIP-Seq (MACS). Genome Biol 9, R137. (doi:10.1186/gb-2008-9-9-r137)

125. Quinlan AR, Hall IM. 2010 BEDTools: a flexible suite of utilities for comparing genomic features. Bioinformatics 26, 841–842. (doi:10.1093/bioinformatics/btq033)

126. Stark, R, Brown G. 2017 DiffBind: Differential Binding Analysis of ChIP-Seq Peak Data. Bioconductor. (doi:10.18129/B9.BIOC.DIFFBIND)

127. McLean CY, Bristor D, Hiller M, Clarke SL, Schaar BT, Lowe CB, Wenger AM, Bejerano G. 2010 GREAT improves functional interpretation of cis-regulatory regions. Nat Biotechnol 28, 495–501. (doi:10.1038/nbt.1630)

128. Bentsen M et al. 2020 ATAC-seq footprinting unravels kinetics of transcription factor binding during zygotic genome activation. Nat Commun 11, 4267. (doi:10.1038/s41467-020-18035-1)

129. Ovek Baydar D et al. 2026 JASPAR 2026: expansion of transcription factor binding profiles and integration of deep learning models. Nucleic Acids Research 54, D184–D193. (doi:10.1093/nar/gkaf1209)

130. Bruse N, Heeringen SJV. 2018 GimmeMotifs: an analysis framework for transcription factor motif analysis. (doi:10.1101/474403)

131. Csárdi G, Nepusz T, Müller K, Horvát S, Traag V, Zanini F, Noom D. 2026 igraph for R: R interface of the igraph library for graph theory and network analysis. (doi:10.5281/ZENODO.7682609)

132. Pérez-Posada A, Lin C-Y, Fan T-P, Lin C-Y, Chen Y-C, Gómez-Skarmeta JL, Yu J-K, Su Y-H, Tena JJ. 2024 Hemichordate cis-regulatory genomics and the gene expression dynamics of deuterostomes. Nat Ecol Evol 8, 2213–2227. (doi:10.1038/s41559-024-02562-x)

133. Thomas Lin Pedersen. In press. ggraph: An Implementation of Grammar of Graphics for Graphs and Networks.

134. Hadley Wickham, Thomas Lin Pedersen, Dana Seidel. 2025 scales: Scale Functions for Visualization.

135. Heinz S et al. 2010 Simple combinations of lineage-determining transcription factors prime cis-regulatory elements required for macrophage and B cell identities. Mol Cell 38, 576–589. (doi:10.1016/j.molcel.2010.05.004)

136. Nechaev S, Fargo DC, Dos Santos G, Liu L, Gao Y, Adelman K. 2010 Global Analysis of Short RNAs Reveals Widespread Promoter-Proximal Stalling and Arrest of Pol II in *Drosophila*. Science 327, 335–338. (doi:10.1126/science.1181421)

