## Supplementary for "Parental care is a genetic capacitor"

for

**Table S1:** Summary of the fitted brms model predicting offspring body size as a function of care treatment, sex, brood size and carcass weight

**Table S2:** Summary of model comparisons between the full model and several alternative models with reduced complexity in the fixed effects or variance structures

**Table S3:** Assembly statistics for three *N. vespilloides* genome assemblies

**Table S4:** BUSCO completeness scores for three *N. vespilloides* genome assemblies

**Table S5:** Annotation statistics for three *N. vespilloides* genome assemblies

**Table S6:** Additional information on gene models [separate .xls file]

**Table S7:** List of marker genes previously experimentally validated and used for cell type annotation, with references [separate .xls file]

**Table S8:** List of marker genes for each cell cluster [separate .xls file]

**Table S9:** Genes encoding soft and rigid cuticle proteins [separate .xls file]

**Table S10:** Differentially expressed genes in Care vs NoCare larvae, detected using bulk and single-nucleus transcriptomes [separate .xls file]

**Table S11:** Mapping statistics for ATAC-seq and CUT&TAG libraries

**Table S12:** Predicted promoters and enhancers and target genes, with information on differential accessibility, differential TF binding and differential activity [separate .xls file]

**Table S13:** Transcription Factors with differential genomic occupancy across Care and No Care larvae as predicted by footprinting [separate .xls file]

**Table S14:** List of identified Hsp83 targets and stalled genes in *D. melanogaster* and their one-to-one ortholog in *N. vespilloides* [separate .xls file]

**Table S15:** Properties of the Care and NoCare gene regulatory networks

**List of Supplementary Figures**

**Figure S1:** Association between brood size and adult body size across care treatments

**Figure S2:** Total phenotypic, among-family, and residual variance estimates from the brms model predicting offspring body size.

**Figure S3:** *N. vespilloides* genome assembly.

**Figure S4:** *N. vespilloides* genomic repeat landscape

**Figure S5:** Quality control metrics for *N. vespilloides* larval head single-nucleus transcriptomes

**Figure S6:** Differentially expressed genes detected across bulk and single-nucleus data

**Figure S7:** Transcription start site (TSS) enrichment and quality control of larval head chromatin accessibility (ATAC-seq) and H3K4me3 histone enrichment (Cut&Tag) data.

**Figure S8:** Alternative peak calling and differential accessibility approaches

**Figure S9:** Additional properties of regulatory changes between Care and No Care groups

**Figure S10:** Additional properties of Care and No Care gene regulatory networks

**Table S1. Summary of the fitted brms model predicting offspring body size as a function of care treatment, sex, brood size and carcass weight.** Fixed effects regression terms are presented as means with associated standard errors and the upper and lower bounds of the 95% credible interval (CI). Residual variation terms are also presented as regression terms as per standard brms output. Among-family terms are shown in standard deviation units.

| Model component | Term | Estimate | SE | CI (lower) | CI (upper) |
|---|---|---|---|---|---|
| **Fixed effects (B)** | (Intercept) | 5.14 | 0.05 | 5.03 | 5.25 |
| | Sex (male) | -0.05 | 0.04 | -0.12 | 0.03 |
| | Care treatment (MC) | -0.29 | 0.07 | -0.42 | -0.16 |
| | Care treatment (NC) | -0.78 | 0.09 | -0.95 | -0.60 |
| | Brood size | -0.02 | 0.01 | -0.04 | -0.01 |
| | Carcass weight | 0.03 | 0.03 | -0.03 | 0.09 |
| | Care treatment (MC) x Brood size | 0.01 | 0.01 | -0.01 | 0.03 |
| | Care treatment (NC) x Brood size | 0.06 | 0.01 | 0.04 | 0.09 |
| **Residual (B)** | Care treatment (FC) | -0.94 | 0.08 | -1.09 | -0.79 |
| | Care treatment (MC) | -0.91 | 0.07 | -1.05 | -0.76 |
| | Care treatment (NC) | -1.29 | 0.08 | -1.45 | -1.13 |
| **Among-family (SD)** | Care treatment (FC) | 0.13 | 0.07 | 0.00 | 0.24 |
| | Care treatment (MC) | 0.06 | 0.05 | 0.00 | 0.15 |
| | Care treatment (NC) | 0.35 | 0.06 | 0.24 | 0.46 |

**Table S2. Summary of model comparisons between the full model (Methods) and several alternative models with reduced complexity in the fixed effects or variance structures.** The difference in the generic expected log-predictive density (ELPD), along with the standard error of the difference, is shown for each model as provided by leave-on-out (loo) comparison.

| Component | Model change | ΔELPD | SE |
|---|---|---|---|
| **Fixed effects** | Exclude treatment x brood size interaction | -2.4 | 2.1 |
| | Exclude treatment x brood size interaction and treatment main effect | -7.7 | 4.2 |
| **Variance components** | Constrain residual variation to be equivalent across treatments | -6.4 | 4.4 |
| | Constrain among-family variation to be equivalent across treatments | -8.2 | 3.3 |
| | Constrain both among-family and residual variances to be equivalent across treatments | -14.3 | 5.8 |

**Table S3. Assembly statistics for three *N. vespilloides* genome assemblies.** The new NVES_3 genome assembly we present in this study is the first chromosome-scale assembly presented for this species. N50 (respectively N90) is the minimum scaffold length such that all scaffolds of this length or longer cover 50% (respectively 90%) of the genome, while L50 is the number of scaffolds reaching this threshold.

| **Metric** | **NVES_3** | **Nves_UGA_2**<br>GCA_048003765.1 | **Nicve_v1**<br>GCF_001412225.1 |
|---|---|---|---|
| Total length (Mb) | 222.9 | 200.0 | 195.3 |
| Number of contigs/scaffolds | 125 | 116 | 4,650 |
| Largest scaffold (Mb) | 40.9 | 16.3 | 1.8 |
| N50 (Mb) | 29.7 | 9.1 | 0.12 |
| L50 | 4 | 8 | 344 |
| N90 (Mb) | 13.2 | 1.4 | 0.02 |
| L90 | 7 | 29 | 1,905 |
| GC content (%) | 32.06 | 31.86 | 31.85 |
| Ns per 100 kbp | 0.4 | 158.5 | 1,648.5 |

**Table S4. BUSCO completeness scores for three *N. vespilloides* genome assemblies.** Completeness of the assembly (Genome) and its annotation (Proteins) was assessed against the endopterygota_odb10 BUSCO dataset, totalling 2,124 genes expected to be single-copy across species.

| **Metric** | **NVES_3** | **Nves_UGA_2** | **Nicve_v1** |
|---|---|---|---|
| | | GCA_048003765 | GCF_001412225.1 |
| **Genome** | | | |
| Complete (%) | 99.4 | 99.4 | 99.2 |
| Single-copy (%) | 97.9 | 97.7 | 97.8 |
| Duplicated (%) | 1.5 | 1.6 | 1.3 |
| Fragmented (%) | 0.1 | 0.1 | 0.4 |
| Missing (%) | 0.5 | 0.5 | 0.5 |
| **Proteins** | | | |
| Complete (%) | 98.6 | 97.1 | 99.3 |
| Single-copy (%) | 97.2 | 95.9 | 97.7 |
| Duplicated (%) | 1.4 | 1.2 | 1.6 |
| Fragmented (%) | 0.7 | 1.3 | 0.4 |
| Missing (%) | 0.7 | 1.6 | 0.3 |

**Table S5. Annotation statistics for three *N. vespilloides* genome assemblies.** Statistics were obtained using the AGAT tool on corresponding annotation files.

| **Metric** | **NVES_3** | **Nves_UGA_2** GCA_048003765.1 | **Nicve_v1** GCF_001412225.1 |
|---|---|---|---|
| Protein-coding genes | 12,848 | 12,424 | 12,642 |
| Total mRNA transcripts | 19,767 | 19,334 | 19,577 |
| Mean transcripts per gene | 1.5 | 1.6 | 1.5 |
| Mean mRNA length (bp) | 10,053 | 10,263 | 10,196 |
| Mean CDS length (bp) | 1,900 | 1,920 | 1,916 |
| Mean exons per mRNA | 6.9 | 7.0 | 7.0 |
| Single-exon mRNAs | 1,023 | 881 | 869 |
| Overlapping gene pairs | 698 | 669 | 715 |

**Table S11.** Mapping statistics for ATAC-seq and CUT&TAG libraries. Total number of reads per library and fraction of reads mapped to the *N. vespilloides* genome.

| Library | Sample | Total Reads | Total Mapped | % Mapped |
|---|---|---|---|---|
| ATAC-seq | A3_Care | 85125999 | 64344095 | 75.00% |
| ATAC-seq | A3_NoCare | 78941916 | 19845203 | 25.00% |
| ATAC-seq | A4_Care | 87556377 | 68060017 | 77.00% |
| ATAC-seq | A4_NoCare | 51569829 | 15549479 | 30.00% |
| ATAC-seq | A5_Care | 86671030 | 74158093 | 85.00% |
| ATAC-seq | A5_NoCare | 90403800 | 71721535 | 79.00% |
| ATAC-seq | A6_Care | 89659185 | 64474050 | 71.00% |
| ATAC-seq | A6_NoCare | 82609135 | 56576258 | 68.00% |
| Cut&Tag | CT1_H3K4me3_Care | 19207135 | 19207135 | 86.36% |
| Cut&Tag | CT1_H3K4me3_NoCare | 17090053 | 17090053 | 51.84% |
| Cut&Tag | CT2_H3K4me3_Care | 11071156 | 11071156 | 76.18% |
| Cut&Tag | CT2_H3K4me3_NoCare | 11378097 | 11378097 | 45.06% |
| Cut&Tag | CT3_H3K4me3_Care | 10181999 | 10181999 | 76.96% |
| Cut&Tag | CT3_H3K4me3_NoCare | 11284550 | 11284550 | 44.45% |
| Cut&Tag | CT4_H3K4me3_Care | 10958539 | 10958539 | 83.76% |

| | | | | |
|---|---|---|---|---|
| Cut&Tag | CT4_H3K4me3_NoCare | 11863115 | 11863115 | 59.21% |

**Table S15. Properties of the Care and NoCare gene regulatory networks.** Number of transcription factors, target genes and connectivity of the networks.

| Metric | Care | NoCare |
|---|---|---|
| Num Genes in Network | 10,691 | 10,621 |
| Num Active Transcription Factors (TFs) | 184 | 182 |
| Mean Connections per TF | 2,778 | 2,808 |
| Median Connections per TF | 2,004 | 1,963 |
| Num Self-Regulated TFs | 47 | 50 |
| Num Target Genes (TG) | 10,507 | 10,439 |
| Mean Connections per TG | 48 | 48 |
| Median Connections per TG | 31 | 32 |
| Mean Emitted Connections | 48 | 48 |
| Mean Received Connections | 48 | 48 |
| Number of common TFs | 180 | 180 |
| Number of exclusive TFs | 4 | 2 |
| Number of common targets | 10,297 | 10,297 |
| Number of exclusive targets | 210 | 142 |

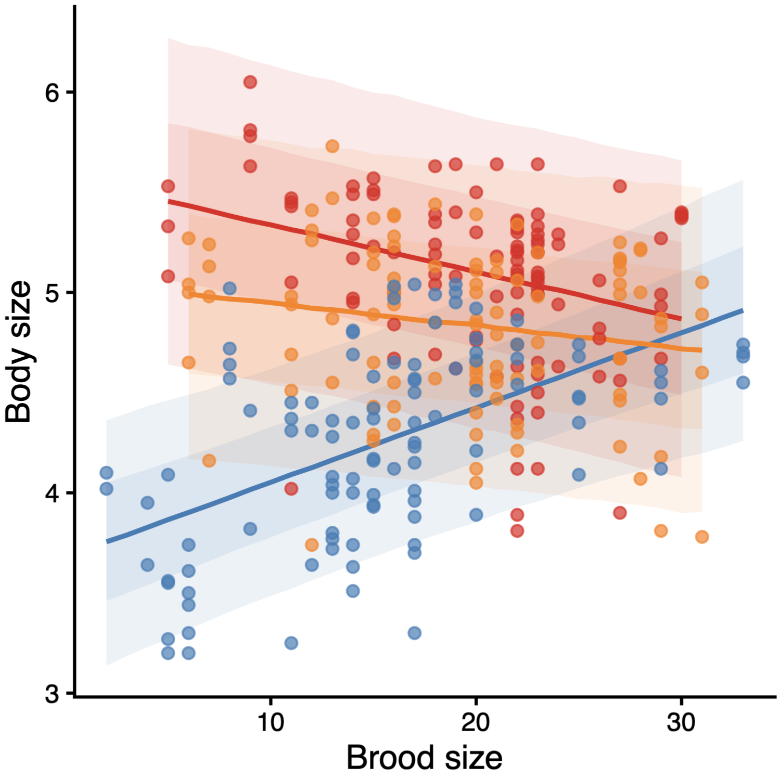

**Figure S1. Association between brood size and adult body size across care treatments.** The effect of brood size on average adult body size differs across treatment groups, with smaller brood size associated with smaller body size in larvae reared without parental care (No Care)NC. Coloured lines and shaded areas show the median with 66% and 95% quantile intervals computed from draws of the posterior predictive distribution, separately for each treatment group. Coloured points show the raw data.

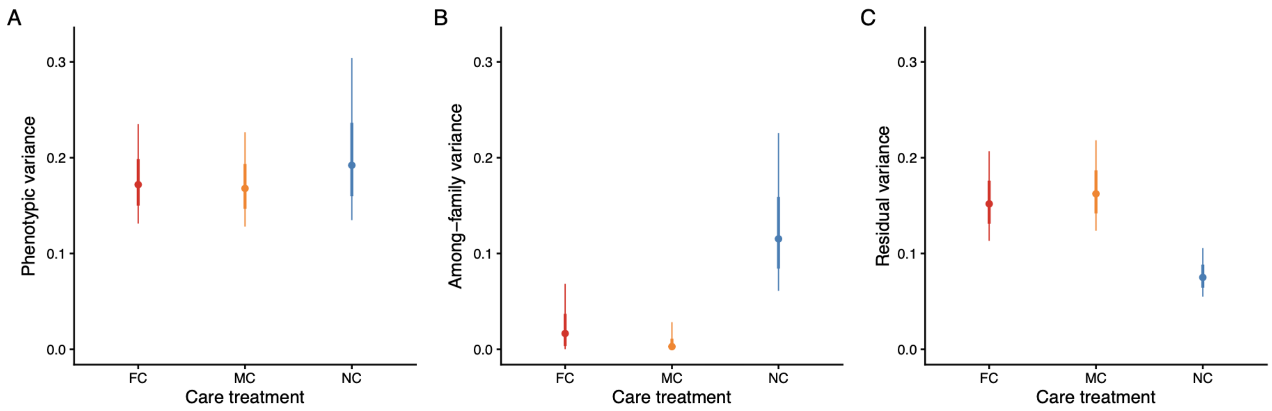

**Figure S2. Estimates from the brms model for among-family (A), residual (B), and phenotypic (C) variances (after accounting for fixed effects within the model).** Within each panel, points and vertical lines show the median with 66% and 95% quantile intervals from draws of the statistical mode separately for each care treatment.

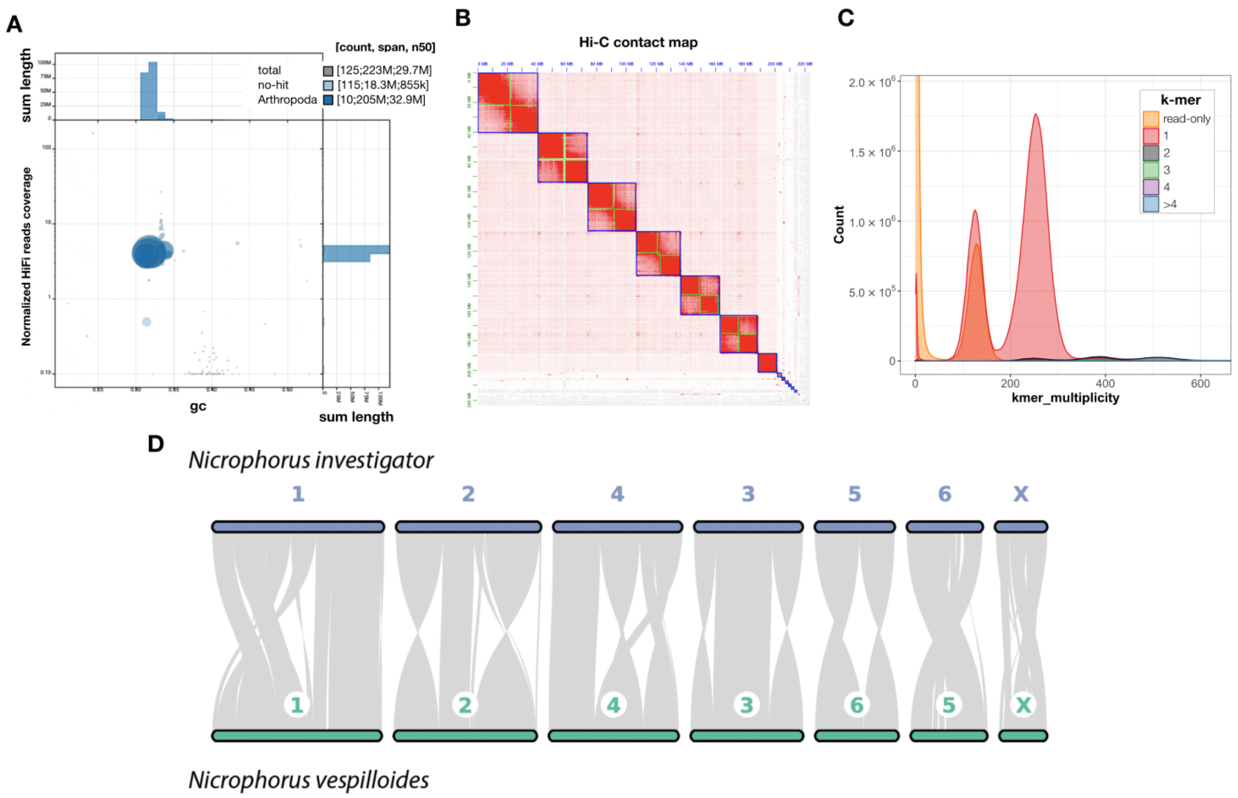

**Figure S3. *N. vespilloides* genome assembly. A.** Coverage-versus-GC content plot generated with BlobTools. Each bubble represents a contig, with size proportional to genomic length and colour indicating taxonomic assignment. Large contigs tightly cluster around a ~35% GC content, show high normalised read coverage and are unambiguously assigned to Arthropoda (totalling 205 Mb), confirming the absence of contamination. **B.** Hi-C contact map showing that the seven chromosome-scale scaffolds (blue boxes) display elevated intra-chromosomal physical contacts, confirming robust scaffolding. **C.** K-mer spectrum of the *N. vespilloides* genome assembly. Low-coverage read-only k-mers (orange curve) correspond to sequencing errors and are absent from the assembly. Half of the k-mers from the heterozygous peak (peak at ~125X coverage) are present in reads only (orange), while the other half are present as single copies in the assembly (red). K-mers from the homozygous peak (~250X) are correctly collapsed and represented as single copies in the assembly. **D.** Synteny conservation between the *Nicrophorus investigator* and *Nicrophorus vespilloides* genomes, demonstrating high collinearity and 1-1 orthology across all seven chromosomes.

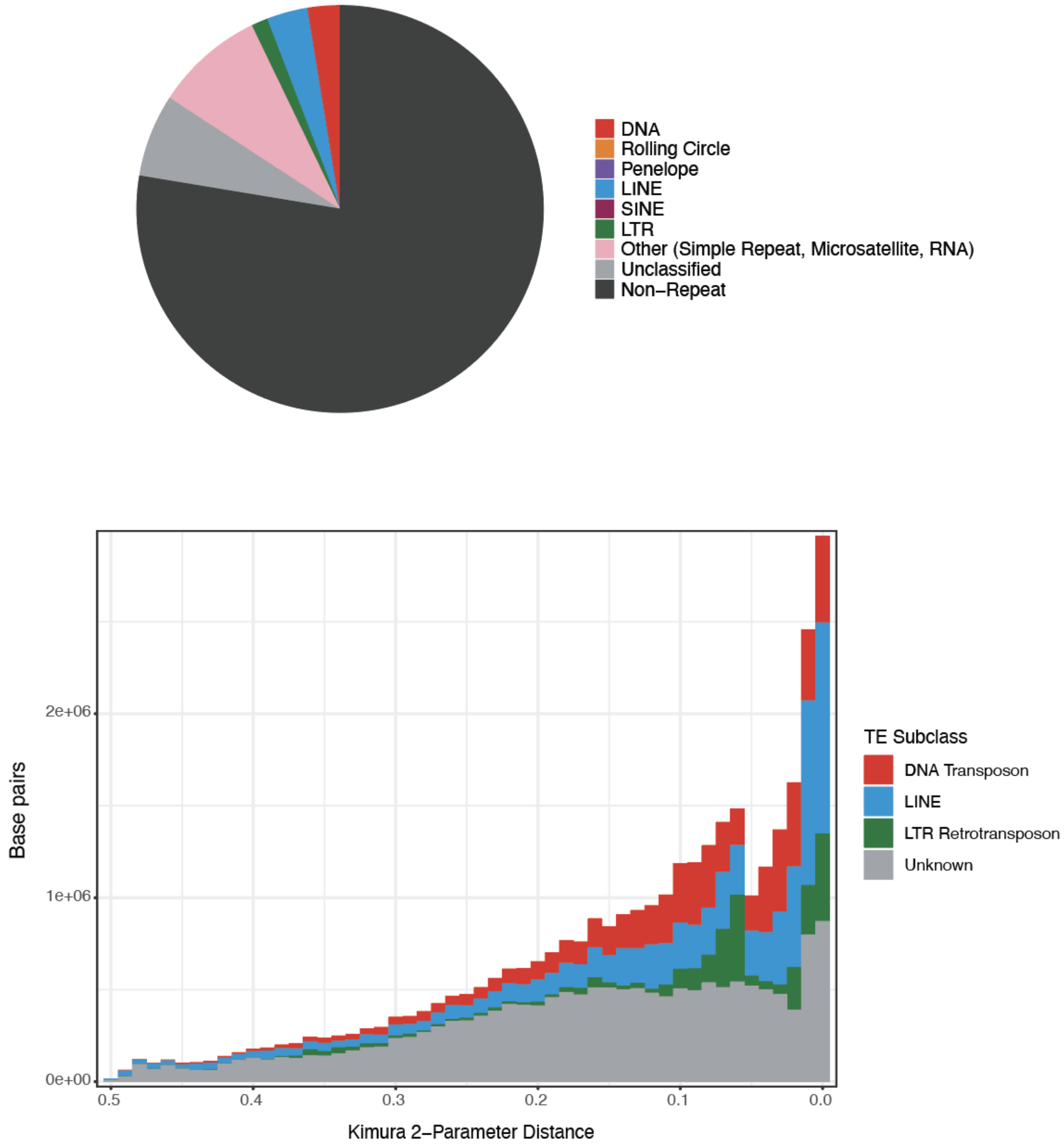

**Figure S4.** ***N. vespilloides*** **genomic repeat landscape. A.** Repetitive element composition. **B.** Repeat landscape showing the genomic coverage (base pairs) of each transposable element (TE) subclass across Kimura 2-parameter distances. Distance from consensus serves as a proxy for relative age of insertion, where a low distance represents recent transposition events.

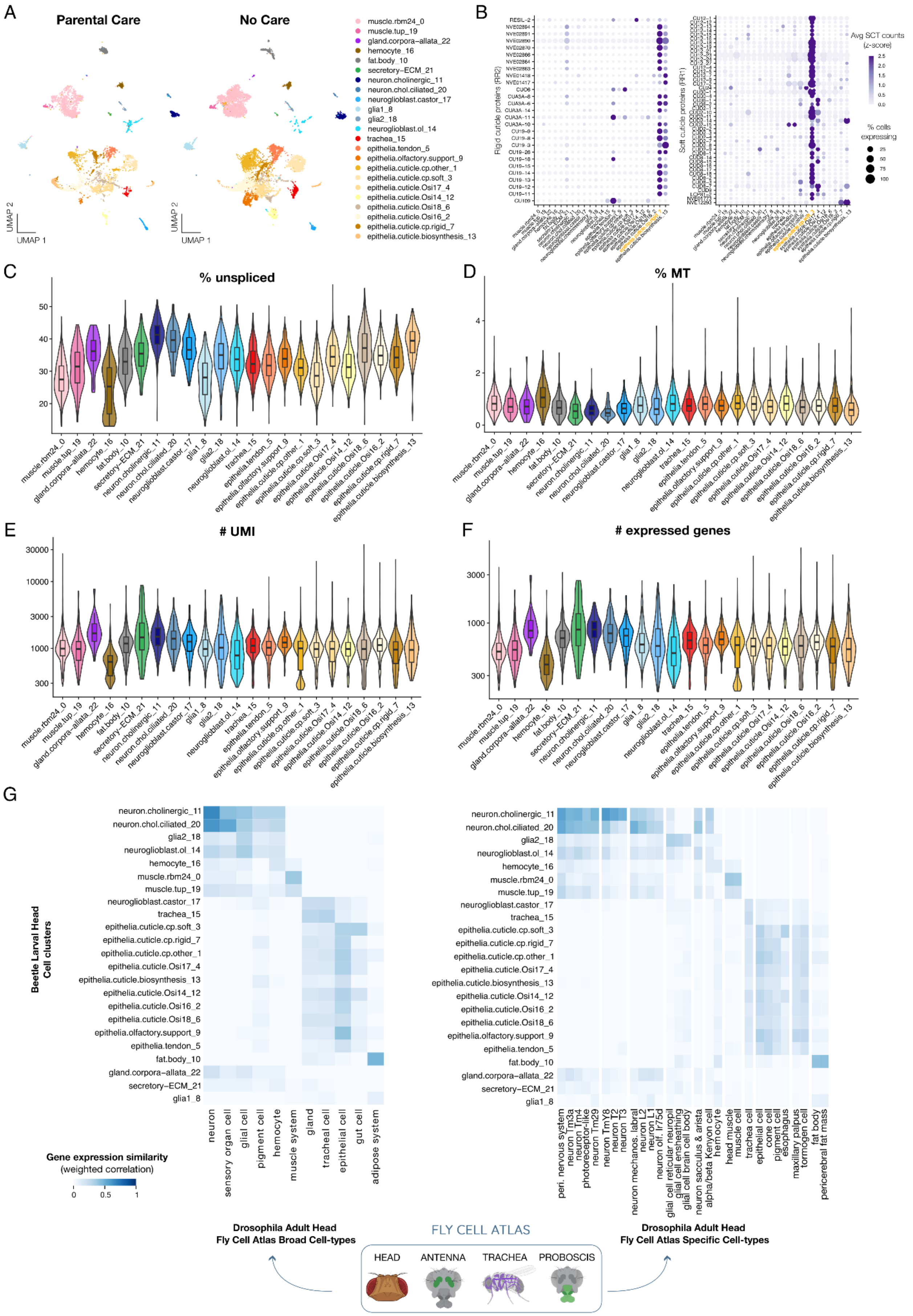
A
Parental Care
No Care
UMAP 1
UMAP 2
muscle.rbm24_0
muscle.tup_19
gland.corpora−allata_22
hemocyte_16
fat.body_10
secretory−ECM_21
neuron.cholinergic_11
neuron.chol.ciliated_20
neuroglioblast.castor_17
glia1_8
glia2_18
neuroglioblast.ol_14
trachea_15
epithelia.tendon_5
epithelia.olfactory.support_9
epithelia.cuticle.cp.other_1
epithelia.cuticle.cp.soft_3
epithelia.cuticle.Osi17_4
epithelia.cuticle.Osi14_12
epithelia.cuticle.Osi18_6
epithelia.cuticle.Osi16_2
epithelia.cuticle.cp.rigid_7
epithelia.cuticle.biosynthesis_13
B
Rigid cuticle proteins (RR2)
Soft cuticle proteins (RR1)
Avg SCT counts (z-score)
% cells expressing
C
% unspliced
D
% MT
E
### UMI
F
### expressed genes
G
Beetle Larval Head Cell clusters
Gene expression similarity (weighted correlation)
neuron
sensory organ cell
glial cell
pigment cell
hemocyte
muscle system
gland
tracheal cell
epithelial cell
gut cell
adipose system
peri. nervous system
neuron Tm3a
neuron Tm4
photoreceptor-like
neuron Tm29
neuron TmY8
neuron T2
neuron T3
neuron mechanos. labral
neuron L2
neuron L1
neuron olf. Ir75d
glial cell reticular neuropil
glial cell ensheathing
glial cell brain cell body
neuron sacculus & arista
alpha/beta Kenyon cell
hemocyte
head muscle
muscle cell
trachea cell
epithelial cell
cone cell
pigment cell
esophagus
maxillary palpus
tormogen cell
fat body
pericerebral fat mass
Drosophila Adult Head
Fly Cell Atlas Broad Cell-types
FLY CELL ATLAS
HEAD
ANTENNA
TRACHEA
PROBOSCIS
Drosophila Adult Head
Fly Cell Atlas Specific Cell-types

**Figure S5. Quality control metrics for *N. vespilloides* larval head single-nucleus transcriptomes. A.** UMAP of all nuclei with annotated cluster, separately for larvae reared with parental care (left) and larvae reared without (right). **B.** Expression of rigid cuticle protein genes (RR2 class, left) and soft cuticle protein genes (RR1 class, right), showing specific expression in rigid.cuticle and soft.cuticle clusters, respectively. **C-F.** Quality control statistics for all retained nuclei across clusters, including: percentage of reads originating from unspliced RNA per nuclei (C), percentage of mitochondrial reads per nuclei (D), number of total unique RNA molecule (UMI) per nuclei (E) and number of expressed genes per nuclei (F). **G.** Gene expression comparisons with reference-annotated *Drosophila* single-cell datasets from the Fly Cell Atlas using Pesci. Expression profiles from our dataset are compared against a merged reference of adult head, antenna, trachea, and proboscis datasets from the Fly Cell Atlas. The panel displays comparisons at the broad cluster level on the left, and with the best-matching specific cell clusters on the right (**Methods**).

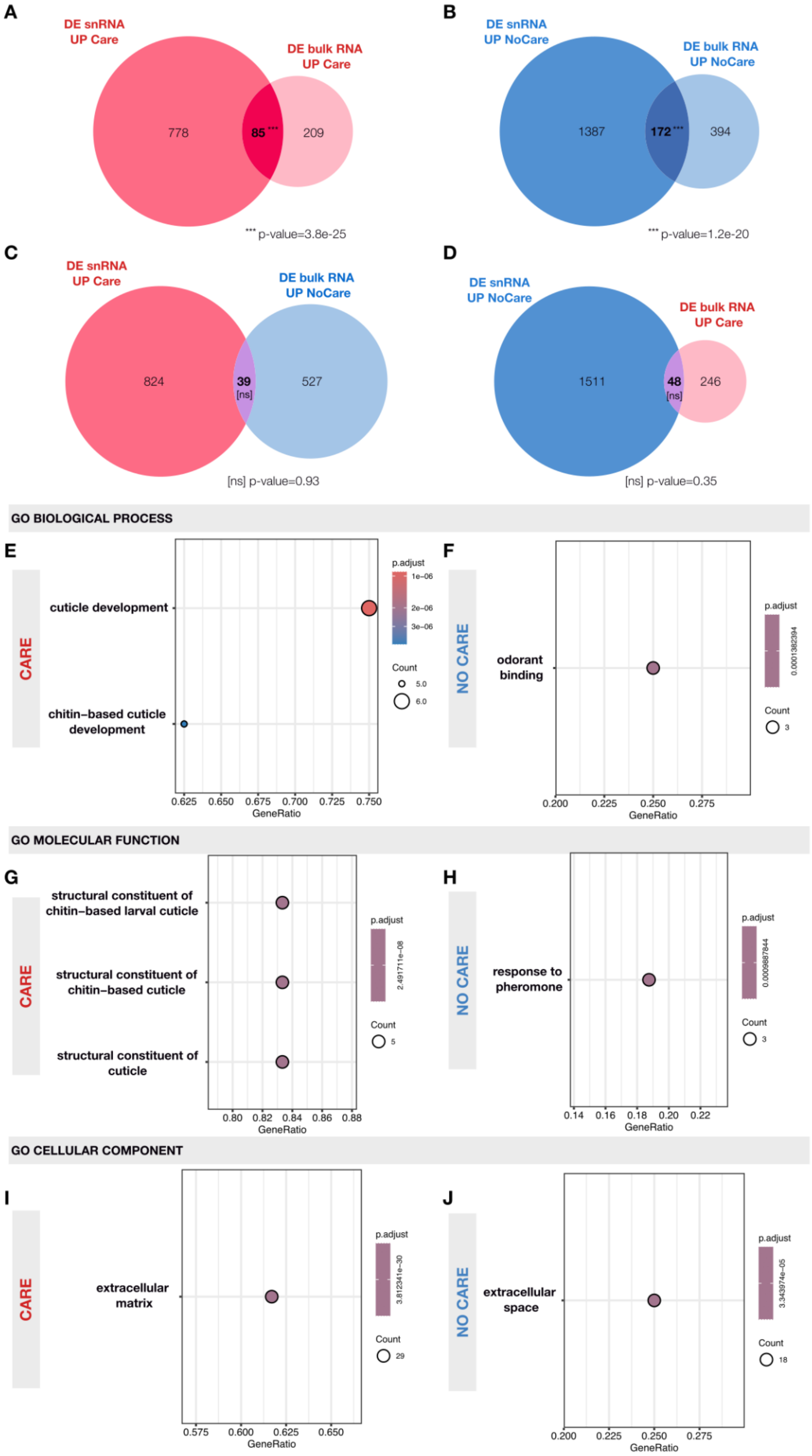

**Figure S6. Differentially expressed genes detected across bulk and single-nucleus data. A–D.** Overlaps between differentially expressed genes detected via snRNA-seq and bulk RNA-seq, with significant overlaps highlighted (hypergeometric *p*-values: * $p < 0.05$, ** $p < 0.01$, *** *p*

< 0.001). Significant overlaps between bulk and snRNA data for Care UP and NoCare UP genes underscore consistency across these approaches, despite their technical and biological differences. **E–J.** Results of Gene Ontology (GO) enrichment tests for genes consistently detected as differentially expressed across both bulk and single-nucleus data (i.e. n=85 consistent Care UP genes in E,G,I and n=172 consistent NoCare UP genes in F,H,J).

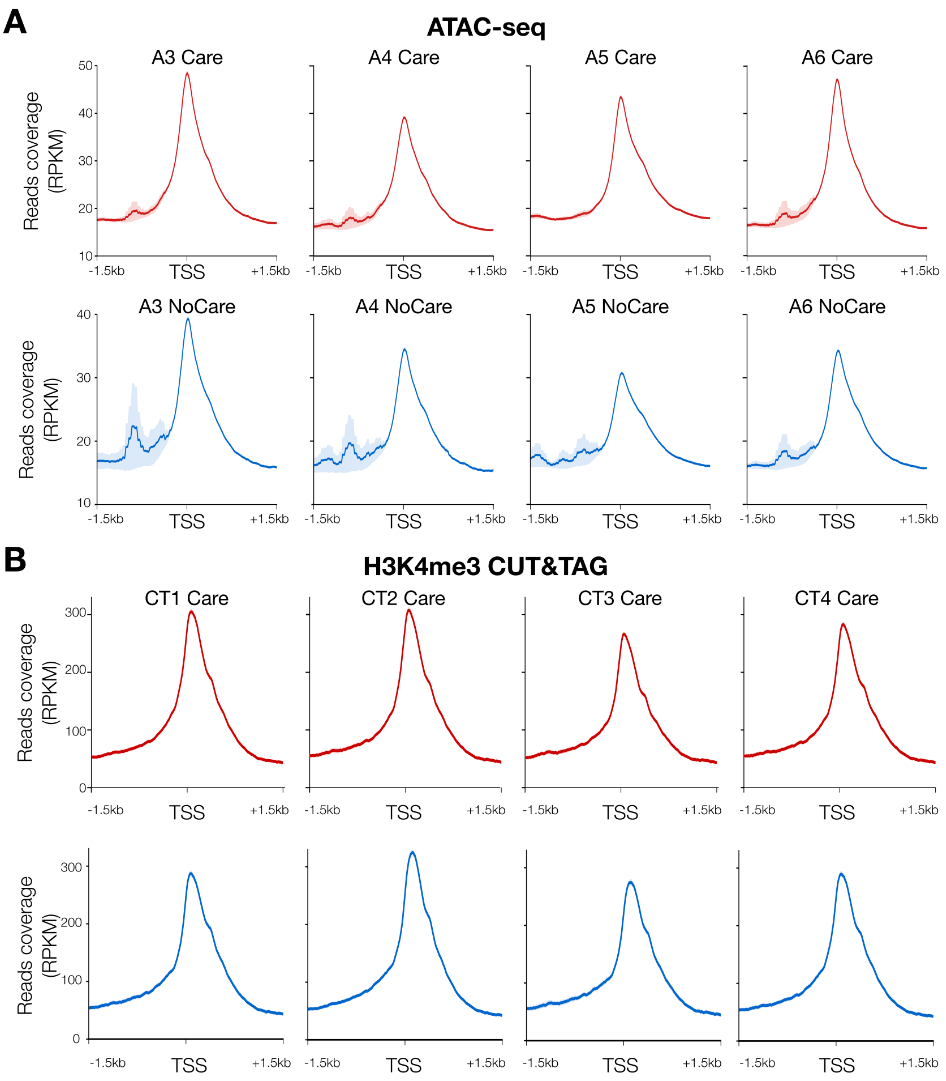

**Figure S7: Transcription start site (TSS) enrichment and quality control of larval head chromatin accessibility (ATAC-seq) and H3K4me3 histone enrichment (Cut&Tag) data. A.** Aggregated ATAC-seq signal relative to annotated transcription start sites (TSS) across the *Nicrophorus vespilloides* genome. The x-axis shows the genomic distance (bp) flanking the TSS, the y-axis normalized coverage (RPKM). **B.** Read coverage enrichment at TSS for H3K4me3 CUT&Tag signal, as in A.

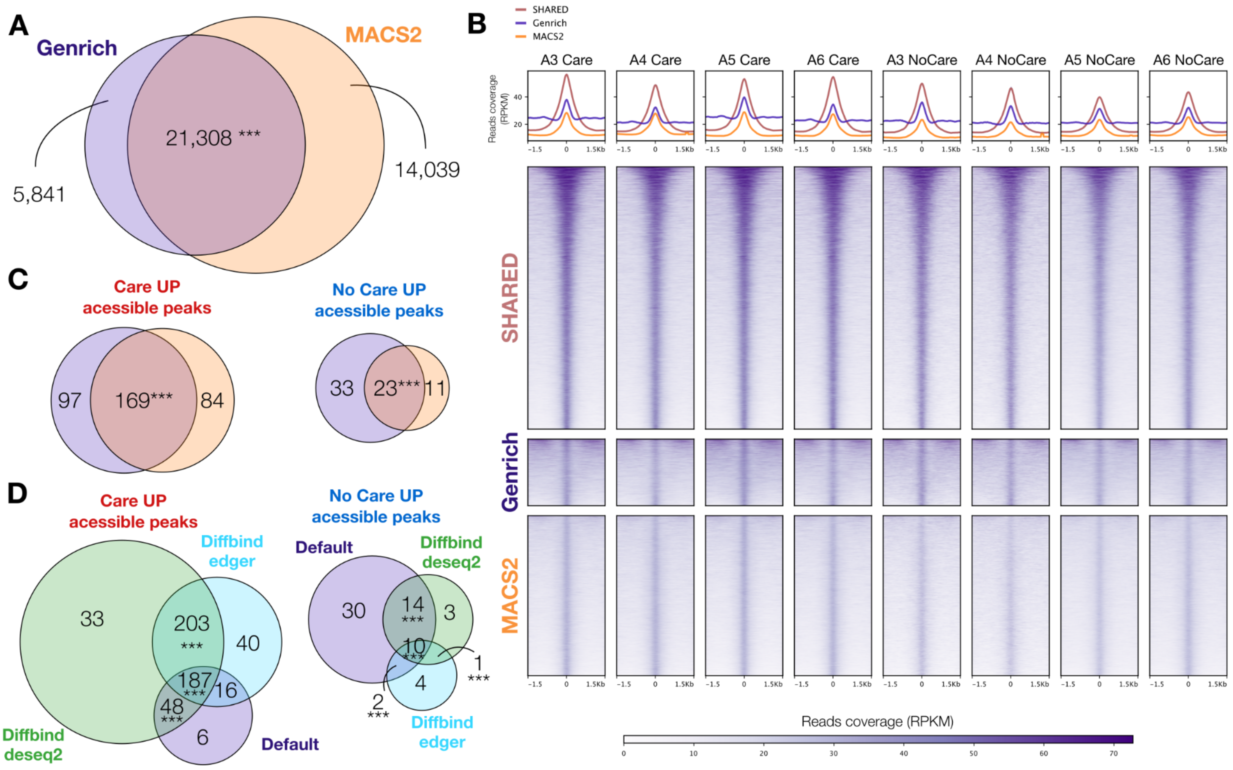

**Figure S8: Alternative peak calling and differential accessibility approaches. A.** Comparison of our Genrich peak set with an alternative approach consisting of calling peaks separately for each replicate with MACS2 and merging reproducible peaks (>= 2 replicates) into a consensus peak set. Both methods recover similar peak set (bedtools Fisher test *p*-values: * $p < 0.05$, ** $p < 0.01$, *** $p < 0.001$). **B.** ATAC-seq read coverage at consensus peak set in each replicate, showing strong enrichment for shared Genrich - MACS2 peaks ('SHARED'). Peaks detected by MACS2 but not Genrich show lower read coverage at peak summit compared to Genrich peaks not detected by MACS2. **C.** Overlap between peaks identified as differentially accessible using the Genrich or MACS2 peak set, *p*-values as in A. Despite MACS2 detecting more peaks, differential accessibility tests do not detect more peaks as differentially accessible. **D.** Overlap between peaks identified as differentially accessible using our Deseq2-based approach (**Methods**) and alternative methods implemented in DiffBind (default parameters, DiffBind edger and DiffBind Deseq2), *p*-values as in A. DiffBind-based methods detect more peaks as differentially accessible due to a less conservative data normalization. DiffBind peaks significantly overlap with our default set and the result of more open chromatin in the Care group is reproduced. DiffBind differential peaks were moreover less correlated with expression shifts, suggesting potential false positives.

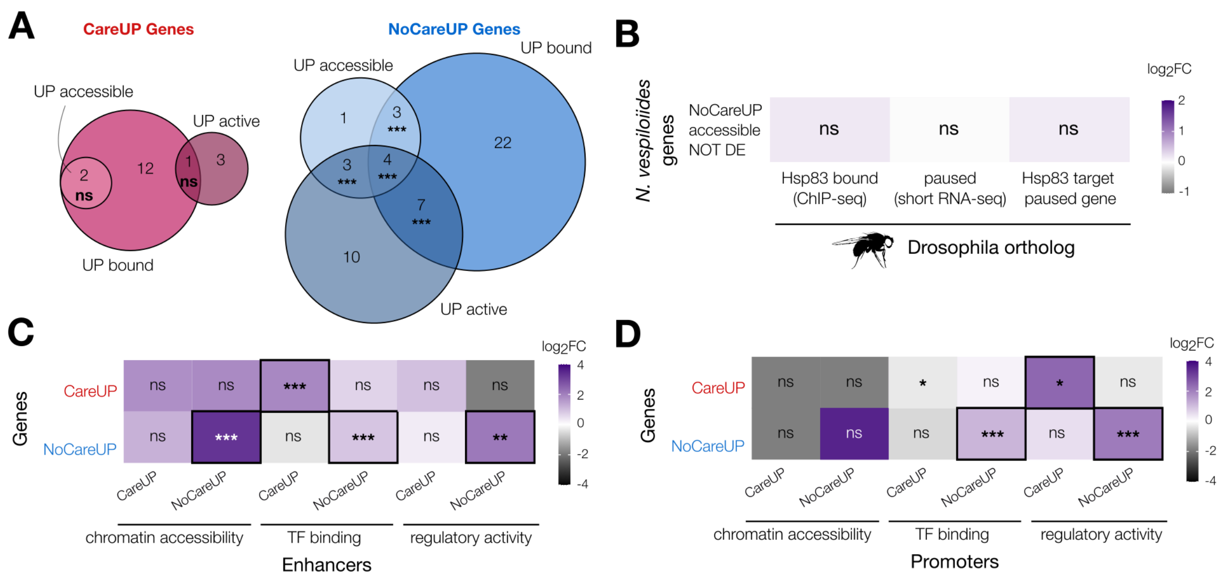

**Figure S9: Additional properties of regulatory changes between Care and No Care groups. A.** Differentially expressed genes (DEG) with concordant changes across regulatory layers. Consistent "No Care" UP DEG show significant overlap (hypergeometric p-values * p<0.05, ** p<0.01, ***p<0.001), indicating concordant changes across examined regulatory layers. **B.** Association tests between *N. vespilloides* genes with higher chromatin accessibility in No Care without differential expression and *D. melanogaster* Hsp83 target and paused genes, using 1-1 orthologs. All hypergeometric p-values were greater than 0.05, indicating a lack of statistically significant association, contrasting with results for Care (**Figure 4g**). Similarly, using Fisher's exact tests did not alter these outcomes, and the resulting odds ratios were systematically smaller for No Care than for Care. **C.** Association tests between differentially expressed genes and differential enhancers (GREAT test hypergeometric p-values * p<0.05, ** p<0.01, ***p<0.001), indicating similar differences between Care and No Care as for the whole set (**Figure 4f**). Enhancers contribute to expression changes mostly through differential TF binding, as expected. **D.** Association tests between differentially expressed genes and differential promoters, as in C. Promoters drive expression changes through change in regulatory activity, as expected.

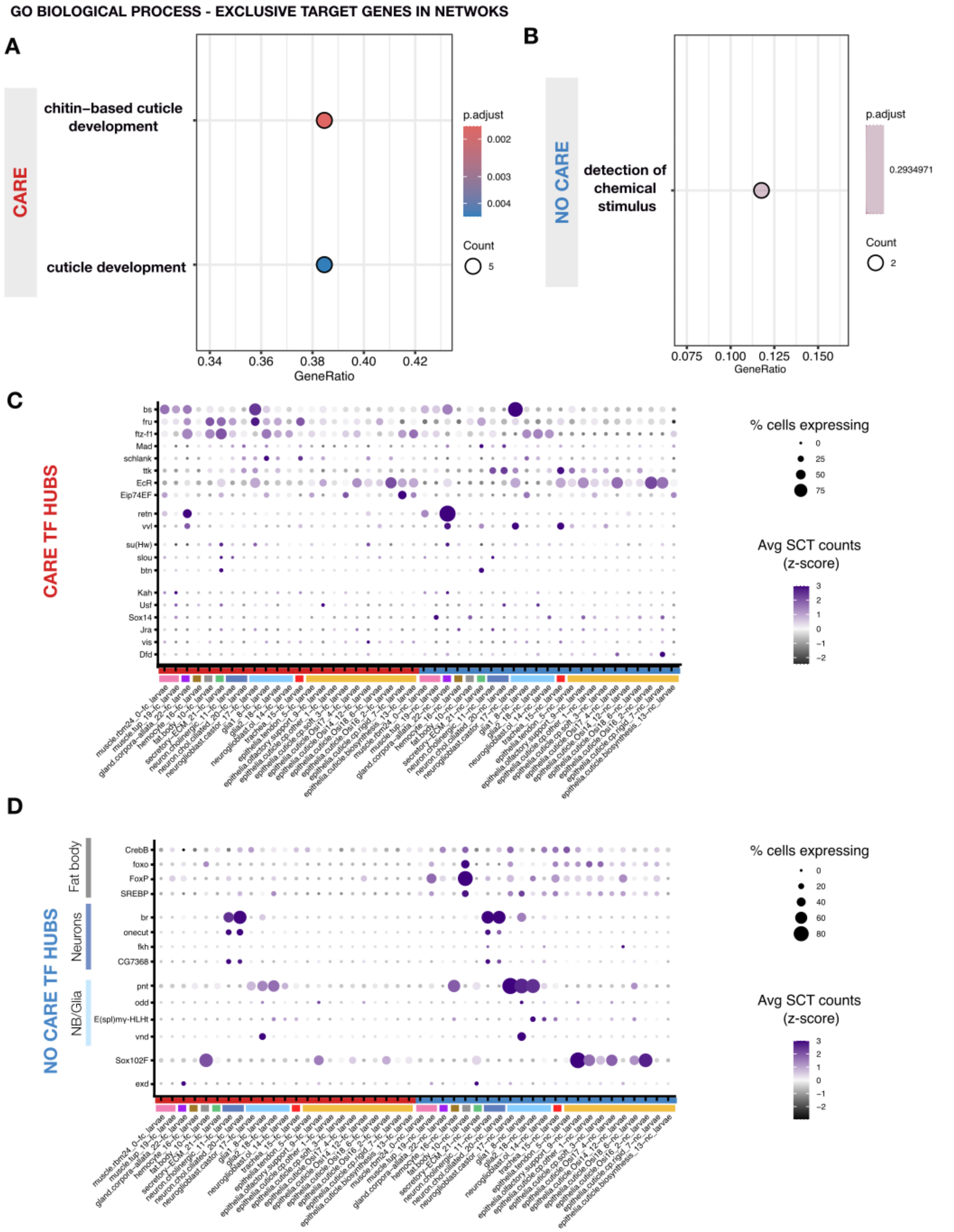

**Figure S10: Additional properties of Care and NoCare gene regulatory networks. A-B.** Results of Gene Ontology (GO) enrichment tests for genes that are exclusive targets of the Care

(**A**) and NoCare (**B**) gene regulatory network. Functional enrichments are similar to enrichment detected for differentially expressed genes (see **Figure S6**). We show the top term for NoCare exclusive targets, despite lack of statistical significance. **C-D.** Cell cluster gene expression for Transcription Factor hubs of the Care (**C**) and NoCare (**D**) regulatory networks. TF hubs of the Care have a globally broader expression, while TF hubs of the No Care network are mostly expressed in fat body cells and neuron-related cell types, consistent with results of differential expression analyses.
